# The Lipidome Landscape of Amiodarone Toxicity: An *in vivo* Lipid-centric Multi-Omics Study

**DOI:** 10.1101/2025.01.21.633980

**Authors:** Nguyen Quang Thu, Jung-Hwa Oh, Nguyen Tran Nam Tien, Se-Myo Park, Nguyen Thi Hai Yen, Nguyen Ky Phat, Tran Minh Hung, Huy Truong Nguyen, Duc Ninh Nguyen, Seokjoo Yoon, Dong Hyun Kim, Nguyen Phuoc Long

## Abstract

Amiodarone is an effective therapy for arrhythmias, its prolonged management may lead to significant adverse drug reactions. Amiodarone-induced hepatoxicity is described by phospholipidosis, hepatic steatosis, cholestatic hepatitis, and cirrhosis. However, the systemic and hepatic lipidome disturbances and underlying toxicological mechanisms remain comprehensively elucidated. Untargeted lipidomics were utilized to analyze serum and liver samples from the rats orally administered a daily dose of amiodarone of either 100 or 300 mg/kg for one week. Changes in the expression of hepatic lipid-related genes were also examined utilizing transcriptomics. We found a higher magnitude of lipidome alterations in the 300 mg/kg than those in the 100 mg/kg groups. Treated animals showed elevated abundances of phosphatidylcholines, ether-linked phosphatidylcholines, sphingomyelins, and ceramides, and decreased levels of triacylglycerols, ether-linked triacylglycerols, and fatty acids. We also found 199 lipid-related differentially expressed hepatic genes between the 300 mg/kg group versus controls, implying lipid metabolism and signaling pathways disturbances. Specifically, elevation of serum phosphatidylcholines and ether-linked phosphatidylcholines, as well as hepatic bismonoacylglycerophosphates were associated with reduced expression of phospholipase genes and elevated expression of glycerophospholipid biosynthesis genes, possibly driving phospholipidosis. Perturbations of sphingolipid metabolism might also be the key events for amiodarone-induced toxicity. Alterations in gene expression levels related to lipid storage and metabolism, mitochondria functions, and energy homeostasis were also found. Collectively, our study characterized the sophisticated perturbations in the lipidome and transcriptome of amiodarone-treated rats and suggested potential mechanisms responsible for amiodarone-induced hepatotoxicity.

## 1. Introduction

Amiodarone is a potent anti-arrhythmic drug used to treat atrial and ventricular arrhythmias by blocking the ion currents. The acute effects of the intravenous (IV) route mainly block the Na^+^ and Ca^2+^ currents, meanwhile the chronic effects of the oral route block the K^+^ currents. The electrophysiological effects of amiodarone lead to the prolongation of the cardiac action potential, and the refractoriness, and prevent/terminate the re-entry [1]. Despite its significant clinical efficacy, amiodarone can cause considerable adverse drug reactions (ADRs) [2, 3]. Amiodarone is highly lipid-soluble, leading to its notable accumulation in adipose tissue and well-perfused organs such as the liver, lungs, and spleen [1]. The loading dose is necessary for the gradual onset of its anti-arrhythmic effects, which might increase the risk of non-cardiac toxicity. In addition, the long-term treatment could be limited due to common drug-drug interactions [4]. Pulmonary toxicity is one of the most serious ADRs of amiodarone, with an estimated about 0.1-17% of cases and mortality rates ranging from 1% to 50% [5]. Patients experiencing pulmonary toxicity commonly exhibit symptoms such as dyspnea, a non-productive cough, malaise, fever, and pleuritic chest pain [3]. Besides, long-term oral amiodarone administration may cause liver injury, which may present as hepatic steatosis, cholestatic hepatitis, or cirrhosis [4, 6]. A meta-analysis evaluated the incidence of liver adverse events in 10,000 person-years between amiodarone and placebo were 54 and 25, respectively [7]. In patients taking amiodarone, liver abnormalities occur in 15-50% of cases, often manifesting as mild increases in serum transaminases (approximately 25%), with approximately 3% developing symptomatic hepatitis [6, 8]. While discontinuing the medication can resolve some cases, others may lead to irreversible liver damage or even death [2, 8]. This underscores a thorough understanding of the underlying molecular mechanisms to guide effective preventive strategies for amiodarone-associated ADRs.

The studies in the *in vitro* and *in vivo* models suggest that impaired mitochondrial function is the mechanism for amiodarone-induced liver injury [9]. The mitochondrial toxicity of amiodarone could lead to an accumulation in ROS and fatty acids (FAs) levels by inhibiting β-oxidation, which may contribute to liver steatosis [10]. Notably, steatosis caused by amiodarone has been reported in human [11]. Importantly, current literature has consistently suggested lipid metabolism-related processes, including oxidative stress, lipid peroxidation, and phospholipidosis, in the context of amiodarone-induced hepatotoxicity [12–14]. However, the effect of amiodarone-induced phospholipidosis has not been well understood. It may be due to the direct binding of the drug to phospholipids, forming indigestible drug–lipid complexes, or inhibiting lysosomal phospholipases [15]. Additional evidence suggests a link between inflammatory, endoplasmic reticulum (ER) stress in the liver, and liver injury associated with amiodarone-induced lipotoxicity [16–18]. Moreover, amiodarone was reported to disrupt cholesterol synthesis by inhibiting DHCR24 - an important enzyme that reduces desmosterol to form cholesterol, resulting in a significant accumulation of desmosterol in cultured cells and serum of treated patients [19]. In the same vein, our recent study that combines transcriptomics and metabolomics revealed an elevation in acylcarnitines (CARs) and disturbances in energy metabolism linked to amiodarone-induced hepatotoxicity, affecting the tricarboxylic acid (TCA) cycle and FA metabolism [20]. Several studies were performed to investigate the alteration of lipid profiles under amiodarone exposure, and the elevation of glycerophospholipids, including phosphatidylcholines (PC), bismonoacylglycerolphosphates (BMP) were reported [21, 22]. However, the alteration of lipidome and the underlying mechanisms related to amiodarone toxicity have not been well-explored and characterized. Collectively, these findings underscore the crucial role of lipid metabolism in the toxic effects of amiodarone and stress the urgent need for a comprehensive understanding of the lipid mechanisms using a lipid-centric omics approach.

Herein, we employed untargeted lipidomics to gain deeper insights into the systemic and local (i.e., hepatic) lipid metabolic changes associated with hepatotoxicity in a well-established rat model orally administrated with amiodarone daily of either 100 mg/kg or 300 mg/kg for one week. We also leveraged liver genome-wide gene expression data to further analyze these lipid metabolic alterations. The dose-dependent lipid metabolic profiles and identified lipid-related genes that regulate complex lipid metabolism were elucidated at the resolution of lipid subclass-dependent changes and the unsaturation of complex lipids. Furthermore, the correlation networks were utilized to illustrate the alteration at the level of lipid species and gene expression. Collectively, our research provided a comprehensive landscape about the alteration of lipid metabolic profiles, as well as discovered and elucidated the potential mechanisms associated with amiodarone-induced liver toxicity.

## 2. Material and methods

### 2.1. Chemicals, reagents, and consumables

Methanol (MeOH), water, isopropanol, acetonitrile, methyl *tert*-butyl ether (MTBE), and toluene at LC-MS grade were purchased from Merck KGaA (Darmstadt, Germany). Ammonium formate and ammonium acetate were obtained from Sigma-Aldrich (St. Louis, MO, USA). The internal standards (IS) of lipids, including SPLASH® LIPIDOMIX® Mass Spec Standard and Deuterated Ceramide LIPIDOMIX® Mass Spec Standard, were provided by Avanti Polar Lipids (Alabaster, AL, USA). The ACQUITY UPLC BEH C18 column (50 mm × 2.1 mm; 1.7 μm) coupled with an ACQUITY UPLC BEH C18 VanGuard pre-column (5 mm × 2.1 mm; 1.7 μm) was purchased from Waters (Milford, MA, USA). Precellys bead tubes for homogenization were obtained from Bertin Technologies (Rockville, MD, USA).

### 2.2. Animals

The current work was the extended analysis using the remaining samples from a previous study [20]. All analyses were conducted according to the Association of Assessment and Accreditation of Laboratory Animal Care International guidelines [20]. The Korea Institute of Toxicology Institutional Review Board approved the experiments (No. 1404-0120). Briefly, male Sprague–Dawley rats were provided from Orient Bio (Seongnam, South Korea) and acclimated for 1 week. Rats were kept in an environment with a temperature of 23 ± 3 °C and relative humidity of 50 ± 10% under a light/dark cycle of 12 h. Rats received amiodarone doses of either 100 mg/kg (n = 10) or 300 mg/kg (n = 10) orally once daily for seven continuous days. The control group (n = 10) was given an equivalent volume of 0.5% (*w/v*) carboxymethyl cellulose (5 mL/kg).

### 2.3. Sample collection and model validation

The collection of serum and tissues followed the established protocol, and the findings of serum biochemical tests and tissue histopathology were previously reported [20]. In summary, a dose-dependent increase of serum phospholipids and total cholesterols was found. Serum aspartate transaminase and alanine transaminase levels were elevated in the 300 mg/kg group. Besides, the blood urea nitrogen levels were significantly increased in rats treated with the amiodarone dose of 300 mg/kg. On the other hand, the reduction of total bilirubin, triacylglycerols (TG), and the average weight of treated groups was also observed. The liver and kidney histopathology revealed no significant changes in the rats treated with amiodarone. Collectively, amiodarone’s one-week treatment-related toxicity was not evident in the rat’s tissue histopathology.

### 2.4. Lipidome extraction for serum

We applied the published protocol for serum lipid extraction with some modifications [23, 24]. Initially, 50 μL of serum was thawed on ice for 30 min and briefly vortexed. Next, 10 μL of a mixture from two ISs solutions (1:1, *v/v*) was spiked to each sample, and the mixtures were vortexed briefly. The sample was then incubated on ice for 20 min, and 300 μL of MeOH (−20 °C) and 1,000 μL of MTBE (−20 °C) were added. The mixtures were vortexed for 10 s and incubated for 20 min at 1,200 *rpm*, 4 °C in a shaker. Then, 250 μL of water was pipetted, followed by vortexing for 20 s. After 10 min of incubation, the mixtures were centrifuged at 14,000 *rcf*, 4 °C for 2 min. Subsequently, 500 μL supernatant of the upper layer was collected and dried at room temperature under N_2_ flow. The dried extracts were stored at −80 °C. Before data acquisition, the dried extracts were resuspended using 200 μL of MeOH:toluene (9:1, *v/v*, 4 °C) and vortex vigorously. The solution was centrifuged for 2 min at 14,000 *rcf*, 4 °C. Then, 150 μL of supernatant was taken into a vial for instrumental analysis, while 20 μL supernatant of each sample was used to make the pooled quality control (QC) sample. The samples were kept in the autosampler at 4 °C during data acquisition.

### 2.5. Lipidome extraction for liver

We followed the previously published protocol with slight adjustments to extract lipids from liver samples [23–25]. First, approximately 30 mg of raw liver tissue was prepared on dry ice. Next, the tissue samples were transferred into a Precellys tube with soft beads on dry ice. Then, 300 μL of MeOH (−80 °C) containing the mixture of two IS solutions (1:1, *v/v*) was added to each tube, followed by two cycles of homogenization by Omni Bead Ruptor Elite homogenizer (Kennesaw, GA, USA) (30 s at 5 m/s for each cycle) with a cooling period on dry ice for 2 min after each cycle. The homogenized tissues were transferred to new 2 mL tubes. Then, 1,000 μL of MTBE at −20 °C was pipetted to each tube and vortexed for 30 s. Next, 20-min incubation was applied for samples (1,500 *rpm* and 4 °C). Subsequently, 250 μL of water (4 °C) was added, vortexed for 20 s, and incubated on ice for 10 min. The extraction mixtures were centrifuged at 16,000 *rcf* for 10 min at 4 °C. Afterward, 500 μL of supernatants from the upper layer were collected and transferred to a new tube, followed by the drying step using the N_2_ flow at room temperature. The dried extracts were stored at −80 °C until analysis. Before LC-MS analysis, dried extracts were reconstituted using MeOH:toluene (9:1, *v/v*, 4 °C) with the final resuspend volume of 300 μL. The mixtures were vortexed briefly, followed by 2-min centrifuged at 14,000 *rcf*, 4 °C. Then, 150 μL of supernatant was transferred into a vial and utilized for LC-MS analysis. Meanwhile, 20 μL was taken to create the pooled QC samples. These samples were stored in the autosampler at 4 °C during analysis.

### 2.6. Lipidomics data acquisition

We utilized the Shimadzu Nexera LC system (Kyoto, Japan) coupled with the X500R Quadrupole Time-of-Flight mass spectrometer (Q-ToF MS), which included a Turbo V™ ion source and Twin Spray probe (SCIEX, Framingham, MA, USA) for the untargeted lipidomics data acquisition [23, 24]. The chromatographic separation was conducted using the ACQUITY UPLC BEH C18 column (50 mm × 2.1 mm; 1.7 μm) connected with an ACQUITY UPLC BEH C18 VanGuard pre-column (5 mm × 2.1 mm; 1.7 μm) [26, 27]. The data-dependent acquisition was applied to acquire the data. The injection volumes were optimized for each type of specimen and each ion mode. For serum, 1.5 μL of each sample was used in positive ion mode (ESI+) and 3 μL in negative ion mode (ESI-). Regarding the liver samples, 1 μL and 2 μL were injected for ESI+ and ESI-, respectively. Gradient elution condition was adapted from the fast LC lipidomics method by Cajka *et al.*, and the MS parameters were followed in our published works [24, 26, 27]. The mass calibration was conducted after every sixth injection using the X500R’s built-in calibrant delivery system.

### 2.7. Lipidomics data processing, annotation, and data treatment

We utilized MS-DIAL version 4.9.0 to process lipidomics raw data, with the parameters followed by our previous works [24]. The MS-DIAL inbuilt library and Fiehn’s lab lipidomics mass spectral library were utilized for lipid annotation, with manual inspection of the experimental and reference MS/MS spectra [28, 29]. Only alignment data of lipids with high confidence of annotation were exported and subsequently processed by MetaboAnalyst 6.0 [30]. First, the features with ≥ 50% missing values were filtered, and the k-nearest neighbors algorithm was applied for missing values imputation. The features with a relative standard deviation of more than 25% in the pooled QC group were also removed. The processed data was subjected to median normalization and log transformation for downstream analysis [24].

### 2.8. Lipidomics data exploration, statistical and functional analyses

The normalized data underwent the Pareto scale before conducting the principal component analysis (PCA) and partial least square-discriminant analysis (PLS-DA). We utilized the five-fold cross-validation procedure and Q^2^ to evaluate the performance of PLS-DA models. Data visualization was conducted using R version 4.2.4 unless stated otherwise [31].

Statistical analysis was performed to determine the significant differential lipids across different doses of amiodarone treatment. We only included subclasses with at least five lipid species to visualize the alteration of lipid subclass abundance. The mixed ion modes data of each specimen were prepared by combining data from two ion modes. The intensity of each subclass was calculated by the sum of intensity from individual species belonging to that subclass. Only the data from one ion mode was kept in the cases of lipid species detected in both ion modes. Then, the one-way analysis of variance coupled with Tukey’s post hoc test was conducted by MetaboAnalyst 6.0, and the cut-off of the false discovery rate (FDR)-adjusted p-value < 0.05 was utilized to consider statistical significance. The heatmaps were plotted by *ComplexHeatmap* package version 2.14.0 [32].

On the other hand, an unpaired two-sided t-test was performed on the level of lipid species and the level of total acyl chain length and total double bonds of lipids in MetaboAnalyst 6.0 and LipidSig 2.0, respectively [30, 33]. Moreover, the lipid-lipid network analysis was conducted to illustrate the alterations of the lipid species in each subclass. The FDR-adjusted p-value cut-off < 0.05 was used to consider statistically differential lipids. The volcano plots of differential lipids were prepared using *EnhancedVolcano* package version 1.16.0. The lipid-lipid network was generated using Gephi. Meanwhile, the plots of alteration trends in total acyl chain length and total double bonds of lipids were illustrated using *ComplexHeatmap* package version 2.14.0 [32]. Furthermore, the mixed ion modes data, including only significant differential lipid species, were subjected to the lipid ontology (LION) enrichment analysis using the module “LION-PCA heatmap.” [34]

### 2.9. Lipid-related gene gene set curation and differential gene expression analysis

The experimental details related to data generation were described in our prior work. In summary, in the 300 mg/kg group and the control group, three liver samples were randomly selected from each group. Subsequently, the total RNA was extracted and the gene expression data were acquired [20]. To investigate the potential mechanisms responsible for the alteration of hepatic lipidome, in this current work, we re-analyzed the transcriptomics data with a focus on other lipid-related genes. The gene expression raw data (.CEL files) preprocessing and quantile normalization were conducted using the *affy* package version 1.76.0 [35]. The normalized data was then submitted to ExpressAnalyst to remove the unannotated genes [36].

We collected the lipid pathways and their corresponding genes from the Kyoto Encyclopedia of Genes and Genomes database, Reactome, and Molecular Signature Database to generate the consensus lipid-related gene set. The gene set was also enriched and validated using the two curated lipid-related gene lists from LipidSig 2.0 [33] and two manually curated lists [37, 38]. A gene was considered a lipid-related gene when it was reported in 2 out of 5 curated sources. Finally, 1056 genes were annotated and subjected to the statistical analysis.

The differential analysis using the *limma* method was conducted to find the significant differentially expressed genes (DEGs). Genes with an FDR < 0.05 were considered statistically significant. Subsequently, the list of DEGs was utilized to generate the bipartite network of lipid-related genes and their corresponding pathways, and the heatmaps of gene expression were generated using the *ComplexHeatmap* package version 2.14.0 [32].

## 3. Results

### 3.1. Amiodarone treatment causes dose-dependent alteration in the systemic and hepatic lipidomes

The PCA results illustrated a tight cluster of the QC samples in both ion modes and specimens **(Supplementary Figure S1)**, indicating the repeatability of the lipidomics data acquisition process and the reliability of the data for subsequent analyses. The PCA scores plots also suggested that the systemic and hepatic lipidomes were altered in the amiodarone-treated groups compared to the control group (**Supplementary Figure S2**). The high-dose groups (300 mg/kg) differed more from the control groups than the low-dose groups (100 mg/kg).

Subsequently, the PLS-DA was performed to differentiate the lipidome among the three groups. The serum samples showed a slight separation between the low-dose and the control groups; meanwhile, a clear separation between the high-dose group and the control group in ESI+ mode (**Figure 1A**). In the ESI-mode, a clear separation between the treated groups and the control group was observed (**Figure 1B**). The cross-validation results demonstrated the good predictive performance of the model in both ion modes (accuracy = 0.767, R^2^ = 0.986, Q^2^ = 0.853 in ESI+; accuracy = 0.833, R^2^ = 0.994, Q^2^ = 0.890 in ESI-) (**Supplementary Figure S3A-B**). Regarding the ESI+ mode data of the liver, we found a clear separation between the high-dose and control groups. However, the lipidome of the low-dose group overlapped with the control group (**Figure 1C**). The liver lipidome of the two treated groups separated from the control group in ESI-mode (**Figure 1D**). The cross-validation results demonstrated weak classification performance in ESI+, but satisfactory classification performance in ESI-(accuracy = 0.797, R^2^ = 0.834, Q^2^ = 0.682 in ESI+; accuracy = 0.763, R^2^ = 0.990, Q^2^ = 0.772 in ESI-, respectively) (**Supplementary Figure S3C-D**).

**Figure 1.**
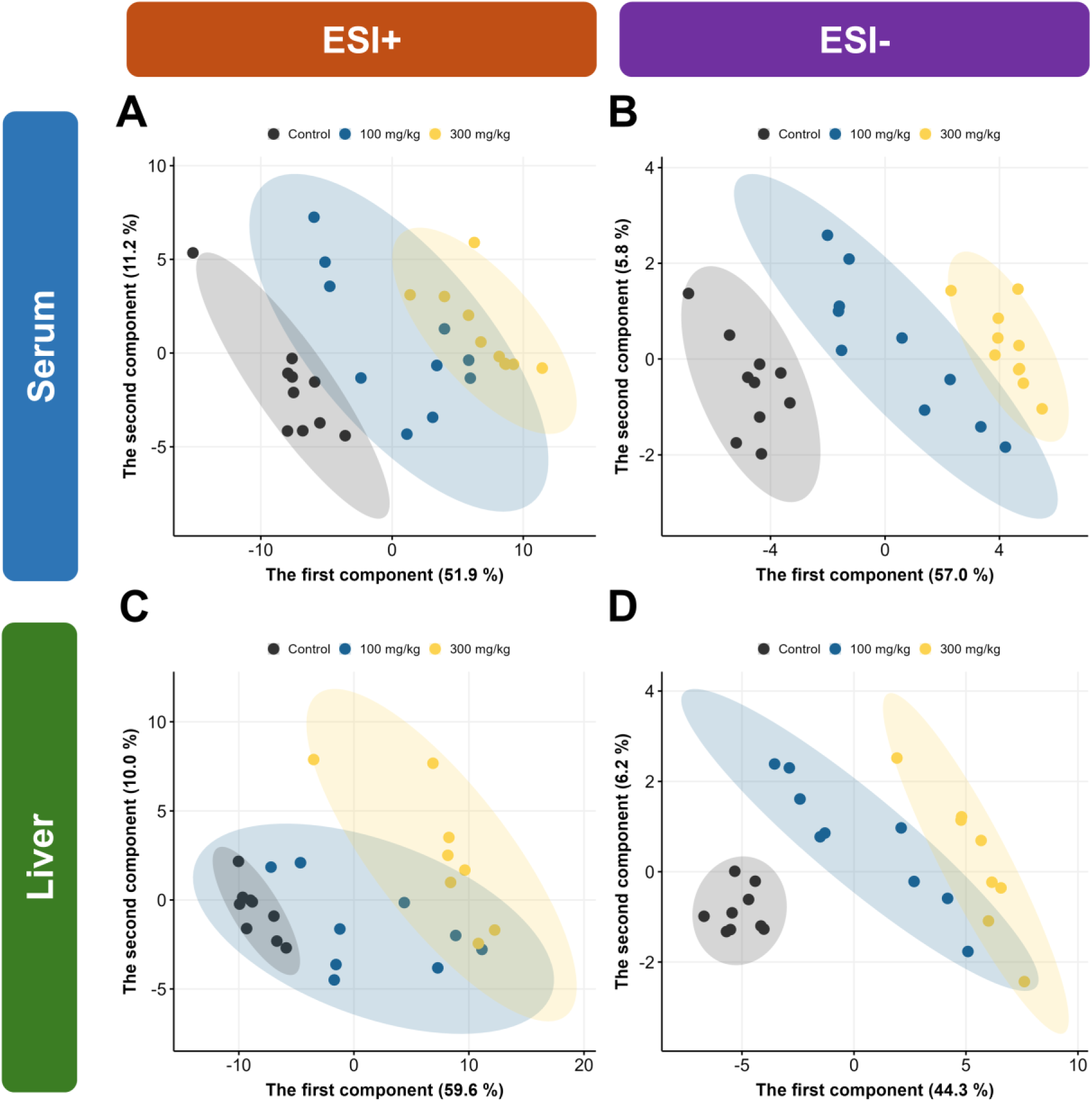
Partial Least Squares-Discriminant Analysis scores plots of serum and liver lipidome of rats treated with amiodarone. (A) Serum lipidome in ESI+ mode. (B) Serum lipidome in ESI-mode. (C) Liver lipidome in ESI+ mode. (D) Liver lipidome in ESI-mode. Abbreviations; ESI+, positive ion mode; ESI-, negative ion mode.

Regardless of biospecimens, we observed the extent of lipid alterations was greater in the 300 mg/kg group compared to the control group than in the 100 mg/kg group relative to the control group. The comparison of the 300 mg/kg group versus the control group revealed the highest number of significant differential lipids compared to other comparisons (100 mg/kg vs. control and 300 mg/kg vs. 100 mg/kg) (**Supplementary Table S1** and **Supplementary Figures S4-5**).

Collectively, exploratory data analysis indicated dose-dependent alterations in the systemic and hepatic lipidome levels of rats exposed to amiodarone at doses of 100 mg/kg and 300 mg/kg.

### 3.2. Amiodarone causes profound systemic lipidome alterations

The serum lipid profiles were significantly altered after one week of amiodarone treatment, and the 300 mg/kg group revealed a higher degree of alteration compared to the 100 mg/kg group. At the level of total acyl chain length and total double bonds, the abundance of lipids with 30-44 carbons increased in treated rats. Meanwhile, the lower abundance of lipids exceeding 48C in the amiodarone-treated groups was determined (**Figure 2A-B**). The alteration of the total amounts of each lipid subclass and the level of lipid species further highlighted these observations. **Supplementary Figure S6A** shows the higher abundance of the lipid subclasses PC, ether-linked phosphatidylcholines [PC(O-)], phosphatidylethanolamines (PE), ether-linked phosphatidylethanolamines [PE(O-)], phosphatidylinositols (PI), ceramides (Cer), sphingomyelins (SM), and cholesteryl esters (CE) in treated groups. Meanwhile, the lower abundances of TGs, ether-linked triacylglycerols [TG(O-)], diacylglycerols (DG), FA, and CAR were observed in the treated rats. When examining individual lipid species, we identified significant elevations in those belonging to PC, PC(O-), SM, and Cer in the treated groups. Conversely, TG and FA were the two main subclasses containing lipids with significantly reduced abundances **(Figure 2C-D, Supplementary Figures S7-8, Supplementary Table S2)**. In addition, the lower total amounts of lipid-containing saturated, monosaturated, and polyunsaturated FA (SFA, MUFA, PUFA) moieties were observed in the amiodarone-treated groups (**Supplementary Figure S6B**).

**Figure 2.**
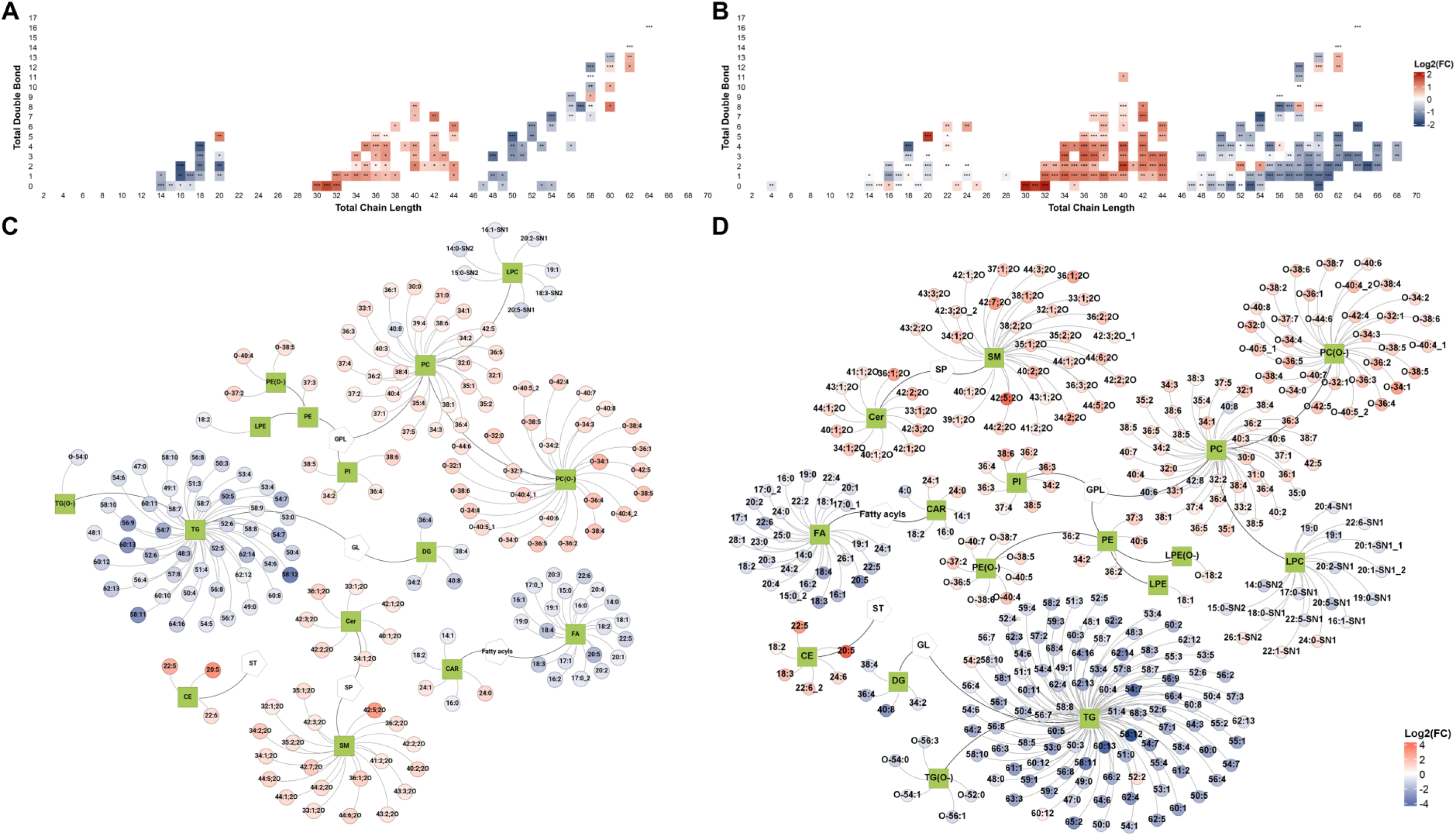
Amiodarone causes profound systemic lipidome alterations. Alteration in the level of total acyl chain length and total double bonds. (A) 100 mg/kg vs. control. (B) 300 mg/kg vs. control. Lipid-lipid network analysis revealed the alteration of significant differential lipids. (C) 100 mg/kg vs. control. (D) 300 mg/kg vs. control. Abbreviations; CAR, acylcarnitines; Cer, ceramides; DG, diacylglycerols; FA, fatty acids; GL, glycerolipids; GPL, glycerophospholipids; LPC, lysophosphatidylcholines; LPE, lysophosphatidylethanolamines; LPE(O-), ether-linked lysophosphatidylethanolamines; PC, phosphatidylcholines; PC(O-), ether-linked phosphatidylcholines; PE, phosphatidylethanolamines; PE(O-), ether-linked phosphatidylethanolamines; PI, phosphatidylinositols; SM, sphingomyelins; SP, sphingolipids; ST, sterol lipids; TG, triacylglycerols; TG(O-), ether-linked triacylglycerols; FDR, false discovery rate; *, FDR < 0.05; **, FDR < 0.01; ***, FDR < 0.001.

### 3.3. Amiodarone causes profound hepatic lipidome alterations

Hepatic lipidome alterations exhibited a similar pattern to systemic lipidome alterations after amiodarone treatment; however, the degree of alteration in lipid subclasses was greater (**Figures 3, Supplementary Tables S3**). The increased abundance of lipids at 30-44C and decreased levels of lipids at 48-62C were found (**Figure 3A-B**). Abundances of phosphatidylglycerols (PG), BMP, PC(O-), PI, PC, cardiolipins (CL), SM, Cer, and CAR were elevated. In contrast, a decrease in the total lipid subclass abundance of TG, TG(O-), and FA was observed (**Supplementary Figure S6C**). We found that PC, PC(O-), and PG were the subclasses that had the highest number of differential lipids with increased abundance in the liver. Moreover, the abundances of all differential lipids belonging to BMP, PG, and CAR were elevated in amiodarone-treated groups, and several had dramatically elevated levels in the liver (**Figure 3C-D, Supplementary Figures S9-10, Supplementary Table S3**). TG and FA were the subclasses with significantly lower abundances after amiodarone exposure (**Figure 3C-D, Supplementary Figures S9-10, Supplementary Table S3**).

**Figure 3.**
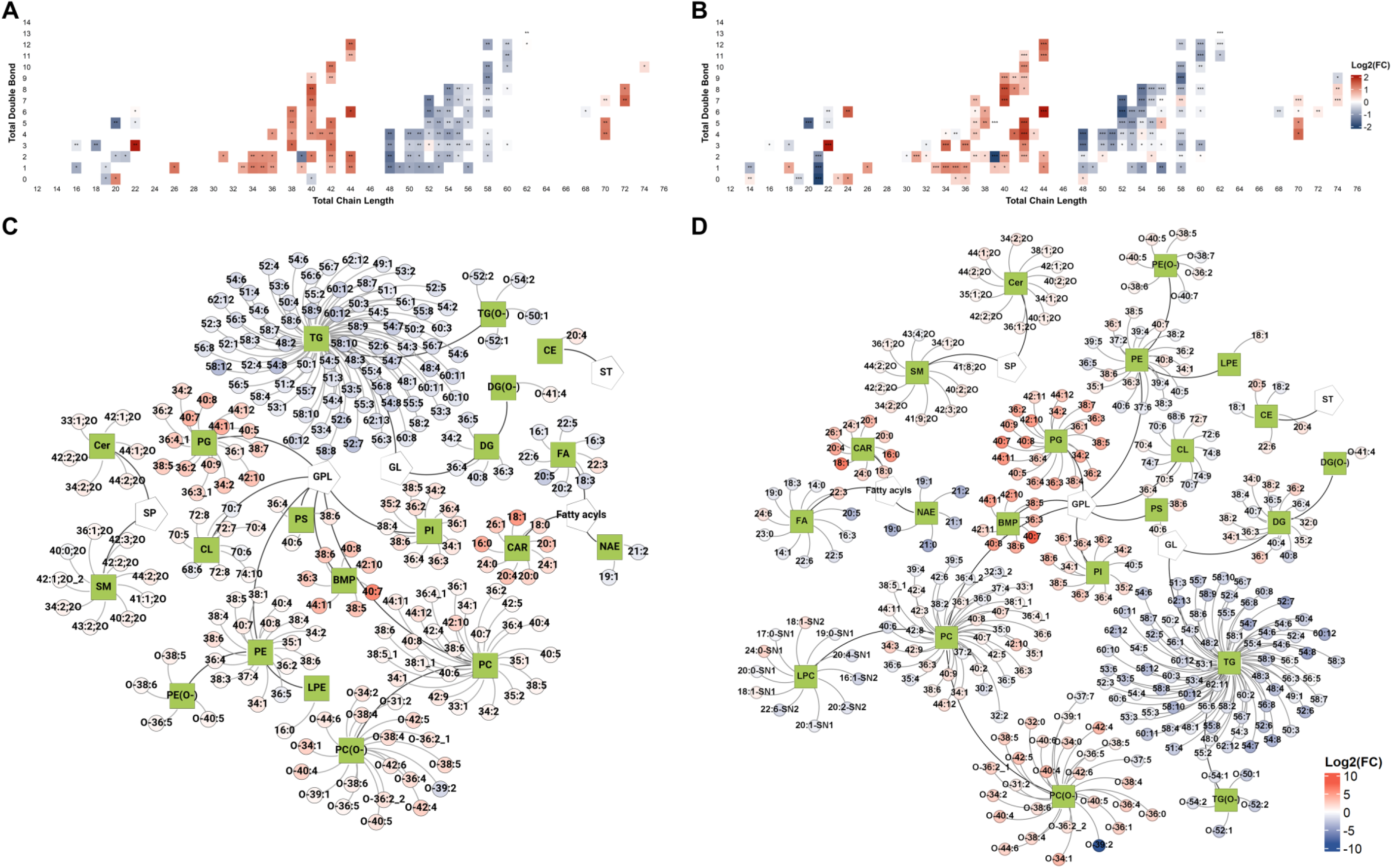
Amiodarone causes profound hepatic lipidome alterations. Alteration in the level of total acyl chain length and total double bonds. (A) 100 mg/kg vs. control. (B) 300 mg/kg vs. control. Lipid-lipid network analysis revealed the alteration of significant differential lipids. (C) 100 mg/kg vs. control. (D) 300 mg/kg vs. control. Abbreviations; BMP, bismonoacylglycerophosphates; CAR, acylcarnitines; CE, cholesteryl esters; Cer, ceramides; CL, cardiolipins; DG, diacylglycerols; DG(O-), ether-linked diacylglycerols; FA, fatty acids; GL, glycerolipids; GPL, glycerophospholipids; LPE, lysophosphatidylethanolamines; NAE, N-acyl ethanolamines; PC(O-), ether-linked phosphatidylcholines; PE, phosphatidylethanolamines; PE(O-), ether-linked phosphatidylethanolamines; PG, phosphatidylglycerols; PI, phosphatidylinositols; PS, phosphatidylserines; SM, sphingomyelins; SP, sphingolipids; ST, sterol lipids; TG, triacylglycerol; TG(O-), ether-linked triacylglycerol; *, FDR < 0.05; **, FDR < 0.01; ***, FDR < 0.001.

We also observed that the lipid droplets and lipid storage in the liver were impaired (**Supplementary Figure S6D**). Notably, the abundances of lipid-containing SFA and PUFA moieties were reduced, including lipids containing C18:2, C18:3, C20:3, C20:5, and C22:5 moieties. Conversely, the elevation of lipid-containing MUFA, particularly lipid species containing C18:1 moiety, was determined in treated groups (**Supplementary Figure S6D**).

Collectively, the lipidomics analysis demonstrated disturbances in the systemic and hepatic lipid metabolism after one week of amiodarone treatment. The high-dose group exhibited a greater extent of lipidome alteration compared to the low-dose group. Abundances of eight lipid subclasses were increased in the treated groups in both serum and liver, including PC, PC(O-), PE, PE(O-), PI, SM, Cer, and CE. Conversely, TG, TG(O-), and FA were three subclasses with lower abundances in both specimens of the treated groups. However, CAR abundances were increased in the liver but decreased in the serum following amiodarone exposure.

### 3.4. Lipid-related gene differential analysis indicated the alteration of underlying lipid metabolism and signaling after one week of amiodarone exposure

One-week amiodarone treatment significantly altered the hepatic lipidome, especially in the high-dose group. Therefore, we investigated the perturbations in hepatic transcriptomic profiles of rats treated with 300 mg/kg of amiodarone (n = 3) compared to the controls (n = 3), to elucidate genes associated with amiodarone-induced lipidome alterations. We found 199 DEGs, including 119 up- and 80 down-regulated genes. **Figure 4** reveals the alteration in the hepatic transcriptome, including DEGs and the corresponding significant pathways after one week of daily 300 mg/kg amiodarone exposure.

**Figure 4.**
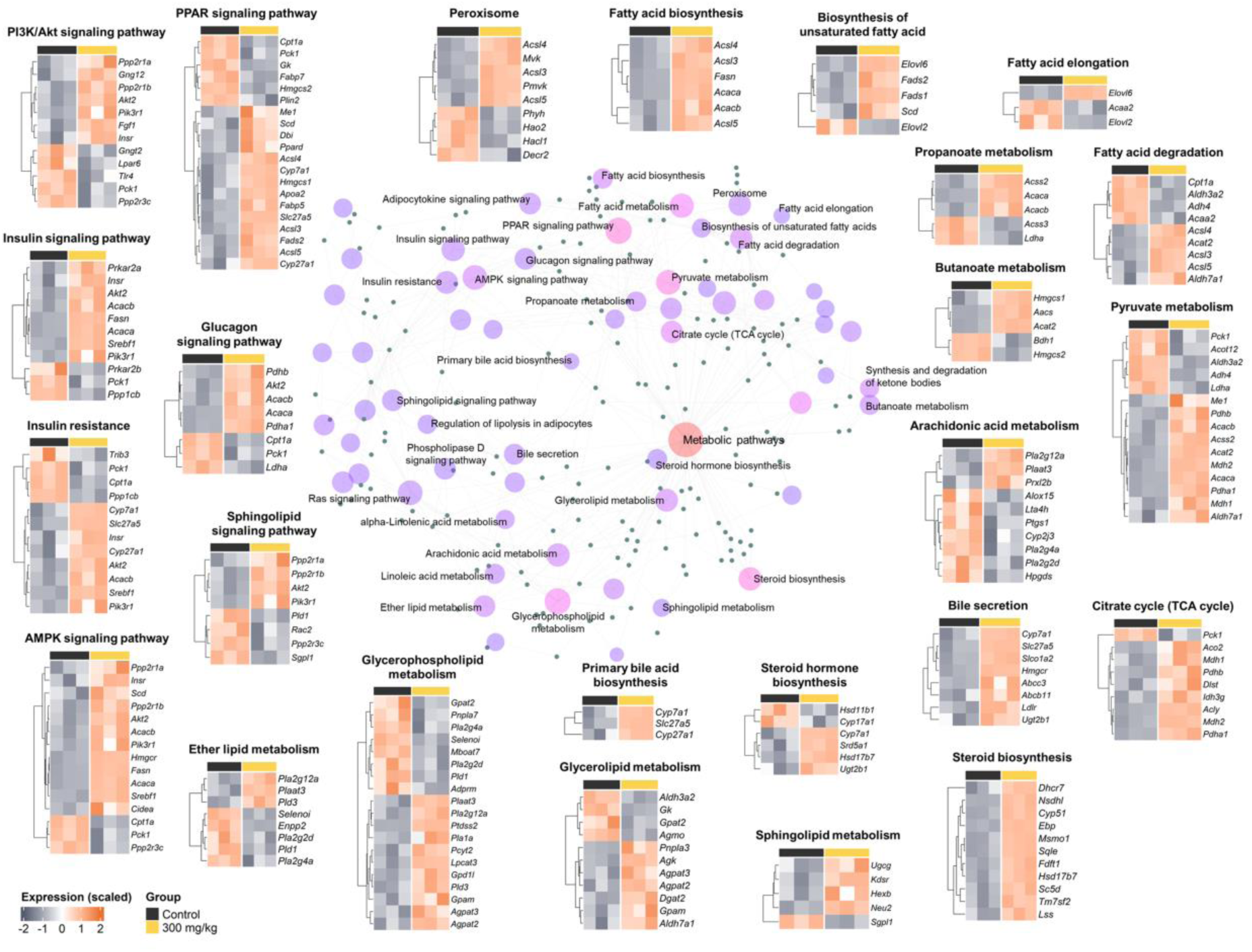
Pathway-level perturbations of lipid-related genes in the rats’ liver after amiodarone treatment. The lipid-lipid-gene network analysis and the heatmaps revealed the alterations in gene expression of DEGs in lipid metabolism and signaling pathways (n = 3 for the 300 mg/kg group, n = 3 for the control group).

Amiodarone exposure significantly affects the hepatic lipidome, including the elevation of glycerophospholipids and sphingolipids levels and the decrease of the glycerolipid level. The transcriptomics analysis results showed perturbations in “glycerophospholipid metabolism,” “sphingolipid metabolism,” and “glycerolipid metabolism” pathways, consistent with the hepatic lipidome alterations. We found that the expression levels of genes encoding phospholipase enzymes (*Pla2g4a*, *Pla2g2d*, and *Pld1*) and other enzymes involved in glycerophospholipid metabolism (*Selenoi*, *Mboat7*, *Pnpla7*, and *Agmo*) were down-regulated.

In contrast, the expression levels of genes involved in glycerophospholipid biosynthesis (*Lpcat3*, *Agpat2*, *Agpat3*, *Gpam*, *Pcyt2*, and *Ptdss2*) were up-regulated. The increased expression levels of *Kdsr*, *Ugcg*, *Hexb*, and *Neu2* in the high-dose treated group, which are involved in the “sphingolipid metabolism” pathway, were also observed. In the “glycerolipid metabolism” pathway, nearly all genes displayed up-regulated expression in the high-dose group compared to the control group. We also found lower expression levels of *Plin2* in the high-dose group, a gene encoded for coated proteins of lipid droplets. In addition, *Npc2*, a gene involved in the transportation process of cholesterol from the endosome to the Golgi apparatus, was suppressed in the 300 mg/kg group compared to the controls.

The expression levels of most genes involved in the mitochondrial function pathways [“Citrate cycle (TCA cycle),” “Pyruvate metabolism,” and “Fatty acid degradation”] were activated. Notably, the genes related to the β-oxidation process in mitochondria (*Cpt1a* and *Acaa2*) and peroxisome (*Decr2*) were down-regulated. In addition, the expression levels of genes involved in the “Fatty acid biosynthesis” pathway, such as *Acly*, *Acaca*, *Acacb*, and *Fasn*, were increased. In the “Biosynthesis of unsaturated fatty acid” pathway, the increased expression levels of *Scd*, *Fads1*, *Fads2*, and *Elovl6* and the decreased expression levels of *Elovl2* were observed in the 300 mg/kg group.

In addition, our results revealed the perturbations in the “Insulin signaling pathway” and “PI3K/Akt signaling pathway,” suggesting amiodarone-induced toxicity might associated with disturbances in insulin regulation in the liver. Moreover, the up-regulation in the expression levels of genes involved in “Steroid biosynthesis,” “Cholesterol metabolism,” “Steroid hormone biosynthesis,” and “Primary bile acid biosynthesis” pathways were found.

## 4. Discussion

Hepatic steatosis, cholestatic hepatitis, and cirrhosis have been reported as ADRs associated with amiodarone, resulting in drug discontinuation in arrhythmia treatment [6]. Understanding the toxicological mechanisms behind these clinical ADRs is crucial for improving the safety of amiodarone use. Our previous study demonstrated the alterations of the systemic and hepatic metabolome and the hepatic transcriptome regarding the FAs and CARs metabolism, as well as in the mitochondria processes under amiodarone exposure [20]. However, the amiodarone-induced lipidome perturbation has not been well-explored and characterized. This study employed lipid-centric multi-omics to shed light on the lipidome alterations and their association with amiodarone toxicity *in vivo*. We found the elevation of systemic and hepatic glycerophospholipid and sphingolipid levels in the treated groups compared to the control group, especially PC, PC(O-), SM, and Cer, or particularly BMP in the liver samples. In addition, the dysregulation in the expression levels of genes regulating the metabolism of these lipids could be key factors of lipotoxicity during amiodarone treatment. We also observed lower levels of both serum and liver TGs and the alterations related to lipid storage and metabolism in the treated groups. Collectively, the putative mechanisms based on hepatic lipidome and transcriptome alterations are summarized in **Figure 5**.

**Figure 5.**
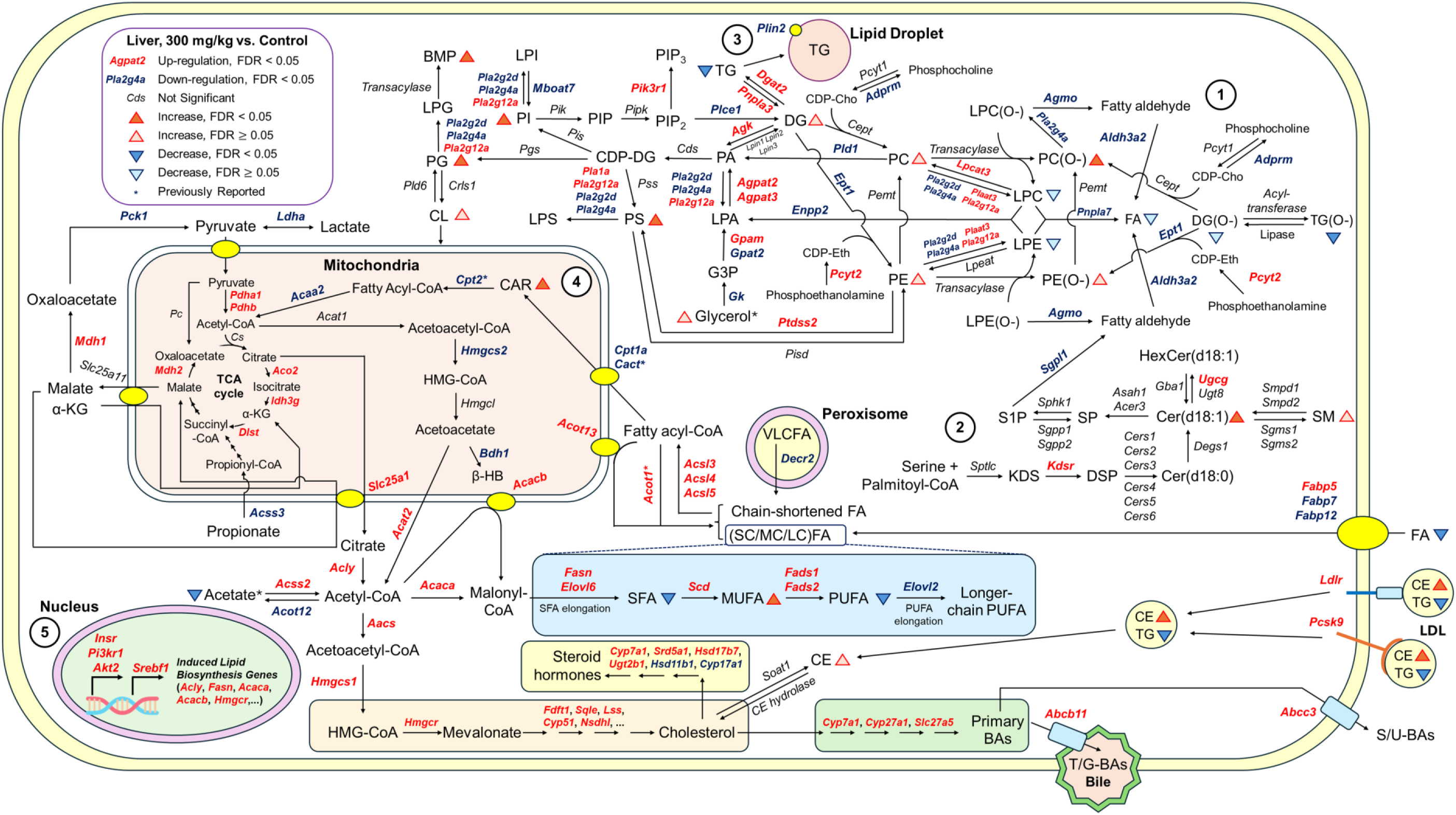
Proposed molecular mechanism of amiodarone-induced hepatotoxicity. Five primary potential mechanisms are illustrated: (1) The alterations of the glycerophospholipid metabolism; (2) The alterations of the sphingolipid metabolism; (3) The alterations of lipid storage and metabolism; (4) Mitochondrial dysfunction and energy homeostasis imbalances; and (5) The activation of lipid-related enzyme genes. Abbreviations; α-KG, α-ketoglutarate; β-HB, β-hydroxybutyrate; BA, bile acids; (S/U)-BAs, sulphation/glucuronidation bile acids; (T/G)-BAs, taurine/glycine-conjugated bile acids; BMP, bismonoacylglycerophosphates; CAR, acylcarnitines; CDP-Cho, cytidine 5’-diphosphocholine; CDP-DG, cytidine 5’-diphosphate diacylglycerols; CDP-Eth, cytidine 5’-diphosphoethanolamine; CE, cholesteryl esters; Cer(d18:0), dihydroceramides; Cer(d18:1), ceramides; CL, cardiolipins; DG, diacylglycerols; DG(O-), ether-linked diacylglycerols; DSP, dihydrosphingosine; FA, fatty acids; (SC/MC/LC/VLC)FA, (short chain/medium chain/long chain/very long chain) fatty acids; (S/MU/PU)FA, (saturated/monosaturated/polysaturated) fatty acids; G3P, glycerol-3-phosphate; HexCer, hexosylceramides; HMG-CoA, 3-hydroxy-3-methylglutaryl coenzyme A; KDS, 3-ketodihydrosphingosine; LPA, lysophosphatidic acids; LPC, lysophosphatidylcholines; LPE, lysophosphatidylethanolamines; LPE(O-), ether-linked lysophosphatidylethanolamines; LPG, lysophosphatidylglycerols; LPI, lysophosphatidylinositols; LPS, lysophosphatidylserines; PA, phosphatidic acids; PC, phosphatidylcholines; PC(O-), ether-linked phosphatidylcholines; PE, phosphatidylethanolamines; PE(O-), ether-linked phosphatidylethanolamines; PG, phosphatidylglycerols; PI, phosphatidylinositols; PIP, phosphatidylinositol monophosphates; PIP_2_, phosphatidylinositol biphosphates; PIP_3_, phosphatidylinositol triphosphates; PS, phosphatidylserines; S1P, sphingosine-1-phosphate; SM, sphingomyelins; SP, sphingosine; TCA, tricarboxylic acid; TG, triacylglycerols; TG(O-), ether-linked triacylglycerols; FDR, false discovery rate.

The elevation of systemic and hepatic glycerophospholipid levels and changes in the expression levels of genes involved in glycerophospholipid metabolism were noted in our study. These observations might be associated with the phospholipidosis status - an ADR during amiodarone treatment [13]. Specifically, the abundances of subclasses PC and PC(O-), among others, showed the highest increase in the treated rats. PC and PC(O-) are two major components of cell membranes and play a crucial role in regulating lipid metabolism, lipoprotein formation, and energy homeostasis [39]. More evidence regarding the perturbations of glycerophospholipid metabolism was obtained from the results of transcriptomics analyses, which revealed decreased expression levels of genes that encode phospholipase enzymes such as *Pla2g4a*, *Pla2g2d*, and *Pld1* in the 300 mg/kg group. The decreased expression levels of phospholipase genes were reported previously in amiodarone-treated rats, which could be the key factor for the higher levels of glycerophospholipids in the treated rats compared to the controls [22]. Conversely, the expression levels of genes related to glycerophospholipid biosynthesis were elevated in the high-dose treated group. It might contribute to the increase in glycerophospholipid levels in amiodarone toxicity. For instance, our study revealed the increased expression levels of *Lpcat3* in the liver of the treated rats. A previous study by Mesens *et al.* showed that amiodarone could activate the expression of the *Lpcat* gene family, which could be associated with higher levels of LPCAT enzymes and result in higher levels of PC and PE in the treated rats [22]. CL is an enriched lipid subclass in the inner membrane of mitochondria, which plays essential roles in bioenergetics, membrane dynamics, and the apoptosis process of mitochondria [40]. Oliveira *et al.* demonstrated the increase of hepatic CLs in rats with nonalcoholic fatty liver disease (NAFLD) and suggested that it could be a regulation effort to maintain energy homeostasis via increased mitochondrial respiratory [41]. Noticeably, amiodarone was also reported to cause NAFLD [42]. Of note in our study, higher amounts of CL were found in the liver of the treated group, which may be associated with the dysregulation of mitochondrial functions induced by amiodarone. In addition, the accumulation of BMPs, mostly species containing acyl chains 22:6, was also found in the liver of treated groups compared to the control group. These lipid species were also reported elsewhere in the *in vitro* and *in vivo* studies and were considered potential biomarkers for amiodarone-induced phospholipidosis [43, 44].

Our study found higher levels of sphingolipid under amiodarone treatment in both serum and liver, particularly SM and Cer, which suggested alterations in the sphingolipid metabolism. SM and Cer play essential roles in cellular membrane dynamics and signaling, and excess sphingolipids abundance cause lipotoxicity in metabolic disorders [45]. The higher expression levels of *Kdsr* were found in the 300 mg/kg group compared to the controls in our study. Of note, KDSR is an enzyme involved in sphingolipids’ *de novo* biosynthesis pathway [46]. Furthermore, the excess of sphingolipids also induces complex effects on glycerophospholipid metabolism [47]. For instance, previous studies in the *in vitro* model showed sphingolipids regulate SREBP1 protein by several mechanisms, including displacing the cholesterol to the cytosol, and cholesterol suppresses the cleavage and activation of SREBP1 [48], or Cer regulates the levels of mature SREBP [49]. Besides, the SREBP1 protein is an important transcription factor that regulates the expression of many genes involved in the biosynthesis of FA, TG, and cholesterol [47].

Lower levels of systemic and hepatic glycerolipids, especially TG, were observed in the treated groups in our study. We also found an alteration in lipid storage and metabolism in the liver, which might be associated with the weight loss noted in rats after one week of amiodarone treatment [20]. In our results, *Plin2* displayed lower gene expression levels in the 300 mg/kg group compared to the controls. Of note, TG and PLIN2 are the primary lipid components and major coated proteins for lipid droplets, respectively [50]. Remarkably, lipid droplets play essential roles in lipid metabolism, energy homeostasis, and cell signaling, especially in protecting against ER stress and mitochondrial damage during autophagy [51].

Previous studies demonstrated amiodarone-induced hepatotoxicity associated with mitochondrial dysfunction and energy homeostasis imbalance [10, 20]. In our study, CAR levels were decreased in serum, meanwhile increased in the liver of the amiodarone-treated rats. Down-regulation of *Cpt1a* gene expression and the accumulation of longer-chain CARs in the liver might suggest the insufficient transportation of FA to the mitochondrial matrix caused by amiodarone [20]. The lower expression levels of *Cpt1a*, *Acaa2*, and *Decr2*, encoding the key enzymes involved in the β-oxidation process, were identified. Notably, DECR2 is an enzyme that is involved in the β-oxidation process in the peroxisome, which helps to shorten the acyl chain of the very long chain FA [52]. Meanwhile, the CPT1A enzyme is involved in the transformation of FA to CAR and transport to the mitochondrial matrix, and the ACAA2 enzyme helps the β-oxidation process occur in mitochondria, thus providing energy and maintaining cellular energy homeostasis [52]. On the other hand, we also found most of the genes belonging to FA metabolism pathways had higher expression levels in the high-dose treated group, which aligned with the previous findings [20]. Of note, in the *in vitro* model using the HepG2 cell line, the activation of key lipogenic genes was observed under amiodarone exposure, including *ACLY*, *FASN*, and *ACACA* [53].

Our study had several limitations that need to be addressed. First, this study investigated rats with daily exposure to toxic doses of amiodarone, but omics profiles were only available at the end of the treatment. Future investigations into the alterations at multiple time points of lipid and transcript profiles should be performed. Next, this research was conducted using serum and liver samples for better elucidation of the underlying mechanism associated with amiodarone-induced hepatoxicity. However, the effects of amiodarone on the lipidome and transcriptome of other organs, which were reported to be susceptible to its toxicity, remain unexplored. Therefore, investigating the lipid and transcript profiles of additional tissue samples, such as the kidney and lung, may provide further insights into the mechanisms underlying amiodarone toxicity at the multi-organ level. On the other hand, this study did not investigate the sex-specific differences of rats in amiodarone toxicity. Subsequent investigations are needed to examine the effects of sex in rats under amiodarone treatment. Furthermore, the toxicity associated with amiodarone when administered intravenously requires further investigation. Collectively, further studies are necessary for a better understanding of the underlying mechanism of amiodarone-induced toxicity. Insights from omics-based *in vivo* toxicity research will help identify potential biomarkers associated with amiodarone toxicity, ultimately improving the safety of this drug in clinical practice.

## 5. Conclusion

In conclusion, our study explored the toxicological effect of a one-week daily amiodarone treatment on both systemic and hepatic lipidomes, as well as hepatic transcriptomes in rats. Our findings indicated significant alterations in systemic and hepatic lipidomes, especially the increase of glycerophospholipid and sphingolipid. Furthermore, lipid storage, lipid metabolism, and mitochondrial function dysregulations were also associated with amiodarone-induced hepatotoxicity. These perturbations might play an important role in amiodarone hepatotoxicity. Our findings revealed the previously underexplored association between amiodarone toxicity and lipid metabolic disturbances, shedding light on the potential mechanisms underlying amiodarone-induced hepatotoxicity.

## CRediT authorship contribution statement

**Nguyen Quang Thu**: Data curation, Methodology, Investigation, Formal analysis, Validation, Writing – Original Draft, Writing – Review & Editing. **Jung-Hwa Oh**: Data curation, Methodology, Validation, Investigation, Formal analysis, Writing – Review & Editing. **Nguyen Tran Nam Tien**: Data curation, Methodology, Software, Formal analysis, Visualization, Validation, Writing – Review & Editing. **Se-Myo Park**: Methodology, Investigation, Validation, Writing – Review & Editing. **Nguyen Thi Hai Yen**: Data curation, Methodology, Validation, Writing – Review & Editing. **Nguyen Ky Phat**: Data curation, Software, Visualization, Writing – Review & Editing. **Tran Minh Hung:** Methodology, Validation, Writing – Review & Editing. **Huy Truong Nguyen**: Methodology, Validation, Writing – Review & Editing. **Duc Ninh Nguyen**: Methodology, Validation, Writing – Review & Editing. **Seokjoo Yoon**: Methodology, Resources, Writing – Review & Editing. **Dong Hyun Kim**: Conceptualization, Methodology, Investigation, Validation, Supervision, Resources, Writing – Review & Editing, Funding acquisition. **Nguyen Phuoc Long**: Conceptualization, Methodology, Investigation, Validation, Supervision, Resources, Writing – Original draft, Writing – Review & Editing, Funding acquisition.

## Declaration of competing interest

The authors stated that the study was carried out in the absence of a commercial or financial relationship that could be interpreted as a potential conflict of interest.

## Fundings

This research was supported by a grant from the Ministry of Food and Drug Safety in 2014 (13182MFDS988) and by the National Research Foundation of Korea (NRF) grant funded by the Korean government (MSIT) (grant No. 2018R1A5A2021242). The funders had no role in study design, data collection, data analysis and interpretation, the manuscript content, or the submission of the manuscript for publication.

## Declaration of generative AI in scientific writing

Grammarly was utilized to verify the grammar and enhance the readability of the manuscript. After using this tool, the authors reviewed and edited the content as needed and took full responsibility for the content of the publication.

## Data availability

The data supporting this study’s findings are available upon reasonable request.

## Appendix A. supplementary data

Supplementary data for this article can be found in the online version.

## Notes

### Competing Interest Statement

The authors have declared no competing interest.

## References

[1] N. Wolkove, M. Baltzan, Amiodarone Pulmonary Toxicity, Can Respir J, 16 (2009) 282540.

[2] J.I. Choi, Y.G. Kim, H.S. Kim, M.N. Kim, J.H. Jeong, H.S. Lee, Y.Y. Choi, J. Shim, Y.H. Kim, Amiodarone is associated with increased mortality compared with other antiarrhythmic drugs in female patients with atrial fibrillation, Eur. Heart J., 44 (2023).

[3] H.-S. You, J.H. Yoon, S.B. Cho, Y.-D. Choi, Y.H. Kim, W. Choi, H.-C. Kang, S.K. Choi, Amiodarone-Induced Multi-Systemic Toxicity Involving the Liver, Lungs, Thyroid, and Eyes: A Case Report, Front. Cardiovasc. Med., 9 (2022).

[4] N. Mujović, D. Dobrev, M. Marinković, V. Russo, T.S. Potpara, The role of amiodarone in contemporary management of complex cardiac arrhythmias, Pharmacol. Res., 151 (2020) 104521.

[5] J. Mankikian, O. Favelle, A. Guillon, L. Guilleminault, B. Cormier, A.P. Jonville-Béra, D. Perrotin, P. Diot, S. Marchand-Adam, Initial characteristics and outcome of hospitalized patients with amiodarone pulmonary toxicity, Respir. Med., 108 (2014) 638–646.

[6] R.M. Colunga Biancatelli, V. Congedo, L. Calvosa, M. Ciacciarelli, A. Polidoro, L. Iuliano, Adverse reactions of Amiodarone, J Geriatr Cardiol., 16 (2019) 552–566.

[7] M. Ruzieh, M.K. Moroi, N.M. Aboujamous, M. Ghahramani, G.V. Naccarelli, J. Mandrola, A.J. Foy, Meta-Analysis Comparing the Relative Risk of Adverse Events for Amiodarone Versus Placebo, Am. J. Cardiol., 124 (2019) 1889–1893.

[8] S. Rankin, D.H. Elder, S. Ogston, J. George, C.C. Lang, A.M. Choy, Population-level incidence and monitoring of adverse drug reactions with long-term amiodarone therapy, Cardiovasc. Ther., 35 (2017) e12258.

[9] S. Nithiyanandam, S. Evan Prince, Toxins mechanism in instigating hepatotoxicity, Toxin Rev., 40 (2021) 616–631.

[10] K. Begriche, J. Massart, M.-A. Robin, A. Borgne-Sanchez, B. Fromenty, Drug-induced toxicity on mitochondria and lipid metabolism: Mechanistic diversity and deleterious consequences for the liver, J. Hepatol., 54 (2011) 773–794.

[11] I.A. González, L.D. Fuller, X. Zhang, D.J. Papke, Jr, L. Zhao, D. Zhang, X. Liao, X. Liu, M.I. Fiel, X. Zhang, Development of a Scoring System to Differentiate Amiodarone-Induced Liver Injury From Alcoholic Steatohepatitis, Am. J. Clin. Pathol., 157 (2021) 434–442.

[12] G. Serviddio, F. Bellanti, A.M. Giudetti, G.V. Gnoni, N. Capitanio, R. Tamborra, A.D. Romano, M. Quinto, M. Blonda, G. Vendemiale, E. Altomare, Mitochondrial oxidative stress and respiratory chain dysfunction account for liver toxicity during amiodarone but not dronedarone administration, Free Radic. Biol. Med., 51 (2011) 2234–2242.

[13] B. Breiden, K. Sandhoff, Emerging mechanisms of drug-induced phospholipidosis, Biological Chemistry, 401 (2020) 31–46.

[14] A. Berson, V. De Beco, P. Lettéron, M.A. Robin, C. Moreau, J. El Kahwaji, N. Verthier, G. Feldmann, B. Fromenty, D. Pessayre, Steatohepatitis-inducing drugs cause mitochondrial dysfunction and lipid peroxidation in rat hepatocytes, Gastroenterol., 114 (1998) 764–774.

[15] N. Anderson, J. Borlak, Drug-induced phospholipidosis, FEBS Lett., 580 (2006) 5533–5540.

[16] N. Erez, E. Hubel, R. Avraham, R. Cohen, S. Fishman, H. Bantel, M. Manns, B. Tirosh, I. Zvibel, O. Shibolet, Hepatic Amiodarone Lipotoxicity Is Ameliorated by Genetic and Pharmacological Inhibition of Endoplasmatic Reticulum Stress, Toxicol. Sci., 159 (2017) 402–412.

[17] E. Hubel, S. Fishman, M. Holopainen, R. Käkelä, O. Shaffer, I. Houri, I. Zvibel, O. Shibolet, Repetitive amiodarone administration causes liver damage via adipose tissue ER stress-dependent lipolysis, leading to hepatotoxic free fatty acid accumulation, Am J Physiol Gastrointest Liver Physiol, 321 (2021) G298–G307.

[18] F. Wandrer, Ž. Frangež, S. Liebig, K. John, F. Vondran, H. Wedemeyer, C. Veltmann, T.J. Pfeffer, O. Shibolet, K. Schulze-Osthoff, H.-U. Simon, H. Bantel, Autophagy alleviates amiodarone-induced hepatotoxicity, Arch Toxicol, 94 (2020) 3527–3539.

[19] P. Simonen, S. Li, N.K. Chua, A.-M. Lampi, V. Piironen, J. Lommi, J. Sinisalo, A.J. Brown, E. Ikonen, H. Gylling, Amiodarone disrupts cholesterol biosynthesis pathway and causes accumulation of circulating desmosterol by inhibiting 24-dehydrocholesterol reductase, J. Intern. Med., 288 (2020) 560–569.

[20] G. Kim, H.-K. Choi, H. Lee, K.-S. Moon, J.H. Oh, J. Lee, J.G. Shin, D.H. Kim, Increased hepatic acylcarnitines after oral administration of amiodarone in rats, J Appl Toxicol., 40 (2020) 1004–1013.

[21] S. Sanoh, Y. Yamachika, Y. Tamura, Y. Kotake, Y. Yoshizane, Y. Ishida, C. Tateno, S. Ohta, Assessment of amiodarone-induced phospholipidosis in chimeric mice with a humanized liver, J. Toxicol. Sci., 42 (2017) 589–596.

[22] N. Mesens, M. Desmidt, G.R. Verheyen, S. Starckx, S. Damsch, R.D. Vries, M. Verhemeldonck, J.V. Gompel, A. Lampo, L. Lammens, Phospholipidosis in Rats Treated with Amiodarone: Serum Biochemistry and Whole Genome Micro-Array Analysis Supporting the Lipid Traffic Jam Hypothesis and the Subsequent Rise of the Biomarker BMP, Toxicol. Pathol., 40 (2012) 491–503.

[23] N.P. Long, N.K. Anh, N.T.H. Yen, N.K. Phat, S. Park, V.T.A. Thu, Y.-S. Cho, J.-G. Shin, J.Y. Oh, D.H. Kim, Comprehensive lipid and lipid-related gene investigations of host immune responses to characterize metabolism-centric biomarkers for pulmonary tuberculosis, Sci. Rep., 12 (2022) 13395.

[24] N.T.H. Yen, N.K. Anh, R.P. Jayanti, N.K. Phat, D.H. Vu, J.-L. Ghim, S. Ahn, J.-G. Shin, J.Y. Oh, N.P. Long, D.H. Kim, Multimodal plasma metabolomics and lipidomics in elucidating metabolic perturbations in tuberculosis patients with concurrent type 2 diabetes, Biochimie, 211 (2023) 153–163.

[25] S.J. Yoon, N.P. Long, K.-H. Jung, H.M. Kim, Y.J. Hong, Z. Fang, S.J. Kim, T.J. Kim, N.H. Anh, S.-S. Hong, S.W. Kwon, Systemic and Local Metabolic Alterations in Sleep-Deprivation-Induced Stress: A Multiplatform Mass-Spectrometry-Based Lipidomics and Metabolomics Approach, J. Proteome Res., 18 (2019) 3295–3304.

[26] T. Cajka, J. Hricko, L. Rudl Kulhava, M. Paucova, M. Novakova, O. Kuda, Optimization of Mobile Phase Modifiers for Fast LC-MS-Based Untargeted Metabolomics and Lipidomics, Int. J. Mol. Sci., 24 (2023) 1987.

[27] N.K. Anh, A. Lee, N.K. Phat, N.T.H. Yen, N.Q. Thu, N.T.N. Tien, H.-S. Kim, T.H. Kim, D.H. Kim, H.-Y. Kim, N. Phuoc Long, Combining metabolomics and machine learning to discover biomarkers for early-stage breast cancer diagnosis, PLOS ONE, 19 (2024) e0311810.

[28] H. Tsugawa, K. Ikeda, M. Takahashi, A. Satoh, Y. Mori, H. Uchino, N. Okahashi, Y. Yamada, I. Tada, P. Bonini, Y. Higashi, Y. Okazaki, Z. Zhou, Z.-J. Zhu, J. Koelmel, T. Cajka, O. Fiehn, K. Saito, M. Arita, M. Arita, A lipidome atlas in MS-DIAL 4, Nat. Biotechnol., 38 (2020) 1159–1163.

[29] T. Kind, K.-H. Liu, D.Y. Lee, B. DeFelice, J.K. Meissen, O. Fiehn, LipidBlast in silico tandem mass spectrometry database for lipid identification, Nat. Methods., 10 (2013) 755–758.

[30] Z. Pang, Y. Lu, G. Zhou, F. Hui, L. Xu, C. Viau, Aliya F. Spigelman, Patrick E. MacDonald, David S. Wishart, S. Li, J. Xia, MetaboAnalyst 6.0: towards a unified platform for metabolomics data processing, analysis and interpretation, Nucleic Acids Research 52 (2024) W398–W406.

[31] R Core Team, R: A Language and Environment for Statistial Computing. R Foundation for Statistical Computing, Vienna, Austria, (2023).

[32] Z. Gu, Complex heatmap visualization, iMeta, 1 (2022) e43.

[33] C.-H. Liu, P.-C. Shen, W.-J. Lin, H.-C. Liu, M.-H. Tsai, T.-Y. Huang, I.-C. Chen, Y.-L. Lai, Y.-D. Wang, M.-C. Hung, W.-C. Cheng, LipidSig 2.0: integrating lipid characteristic insights into advanced lipidomics data analysis, Nucleic Acids Res., 52 (2024) W390–W397.

[34] M.R. Molenaar, A. Jeucken, T.A. Wassenaar, C.H.A. van de Lest, J.F. Brouwers, J.B. Helms, LION/web: a web-based ontology enrichment tool for lipidomic data analysis, Gigascience, 8 (2019).

[35] L. Gautier, L. Cope, B.M. Bolstad, R.A. Irizarry, affy—analysis of Affymetrix GeneChip data at the probe level, Bioinformatics, 20 (2004) 307–315.

[36] P. Liu, J. Ewald, Z. Pang, E. Legrand, Y.S. Jeon, J. Sangiovanni, O. Hacariz, G. Zhou, J.A. Head, N. Basu, J. Xia, ExpressAnalyst: A unified platform for RNA-sequencing analysis in non-model species, Nature Communications, 14 (2023) 2995.

[37] M. Xu, Y.-y. Guo, D. Li, X.-f. Cen, H.-l. Qiu, Y.-l. Ma, S.-h. Huang, Q.-z. Tang, Screening of Lipid Metabolism-Related Gene Diagnostic Signature for Patients With Dilated Cardiomyopathy, Front. Cardiovasc. Med., 9 (2022).

[38] J. Li, Q. Li, Z. Su, Q. Sun, Y. Zhao, T. Feng, J. Jiang, F. Zhang, H. Ma, Lipid metabolism gene-wide profile and survival signature of lung adenocarcinoma, Lipids Health Dis., 19 (2020) 222.

[39] J.N. van der Veen, J.P. Kennelly, S. Wan, J.E. Vance, D.E. Vance, R.L. Jacobs, The critical role of phosphatidylcholine and phosphatidylethanolamine metabolism in health and disease, Biochim. Biophys. Acta - Biomembr., 1859 (2017) 1558–1572.

[40] R.H. Houtkooper, F.M. Vaz, Cardiolipin, the heart of mitochondrial metabolism, Cell. Mol. Life Sci., 65 (2008) 2493–2506.

[41] D.T. Oliveira, A.B. Chaves-Filho, M.Y. Yoshinaga, N.C.N. Paiva, C.M. Carneiro, S. Miyamoto, W.T. Festuccia, R. Guerra-Sá, Liver lipidome signature and metabolic pathways in nonalcoholic fatty liver disease induced by a high-sugar diet, J Nutr Biochem, 87 (2021) 108519.

[42] L.G. Di Pasqua, M. Cagna, C. Berardo, M. Vairetti, A. Ferrigno, Detailed Molecular Mechanisms Involved in Drug-Induced Non-Alcoholic Fatty Liver Disease and Non-Alcoholic Steatohepatitis: An Update, Biomedicines, 2022.

[43] M. Rabia, V. Leuzy, C. Soulage, A. Durand, B. Fourmaux, E. Errazuriz-Cerda, R. Köffel, A. Draeger, P. Colosetti, A. Jalabert, M. Di Filippo, A. Villard-Garon, C. Bergerot, C. Luquain-Costaz, P. Moulin, S. Rome, I. Delton, F. Hullin-Matsuda, Bis(monoacylglycero)phosphate, a new lipid signature of endosome-derived extracellular vesicles, Biochimie, 178 (2020) 26–38.

[44] K.L. Thompson, J. Zhang, S. Stewart, B.A. Rosenzweig, K. Shea, D. Mans, T. Colatsky, Comparison of urinary and serum levels of di-22:6-bis(monoacylglycerol)phosphate as noninvasive biomarkers of phospholipidosis in rats, Toxicol. Lett., 213 (2012) 285–291.

[45] B. Chaurasia, S.A. Summers, Ceramides – Lipotoxic Inducers of Metabolic Disorders, TEM, 26 (2015) 538–550.

[46] M.E. Spears, N. Lee, S. Hwang, S.J. Park, A.E. Carlisle, R. Li, M.B. Doshi, A.M. Armando, J. Gao, K. Simin, L.J. Zhu, P.L. Greer, O. Quehenberger, E.M. Torres, D. Kim, De novo sphingolipid biosynthesis necessitates detoxification in cancer cells, Cell Rep., 40 (2022) 111415.

[47] S. Rodriguez-Cuenca, V. Pellegrinelli, M. Campbell, M. Oresic, A. Vidal-Puig, Sphingolipids and glycerophospholipids – The “ying and yang” of lipotoxicity in metabolic diseases, Prog. Lipid Res., 66 (2017) 14–29.

[48] S. Scheek, M.S. Brown, J.L. Goldstein, Sphingomyelin depletion in cultured cells blocks proteolysis of sterol regulatory element binding proteins at site 1, Proc. Natl. Acad. Sci. U.S.A., 94 (1997) 11179–11183.

[49] T.S. Worgall, R.A. Juliano, T. Seo, R.J. Deckelbaum, Ceramide Synthesis Correlates with the Posttranscriptional Regulation of the Sterol-Regulatory Element-Binding Protein, Arterioscler Thromb Vasc Biol., 24 (2004) 943–948.

[50] H. Itabe, T. Yamaguchi, S. Nimura, N. Sasabe, Perilipins: a diversity of intracellular lipid droplet proteins, Lipids Health Dis, 16 (2017) 83.

[51] J.A. Olzmann, P. Carvalho, Dynamics and functions of lipid droplets, Nat. Rev. Mol. Cell Biol., 20 (2019) 137–155.

[52] S.R. Nagarajan, L.M. Butler, A.J. Hoy, The diversity and breadth of cancer cell fatty acid metabolism, Cancer & Metabolism, 9 (2021) 2.

[53] J. Allard, S. Bucher, J. Massart, P.-J. Ferron, D. Le Guillou, R. Loyant, Y. Daniel, Y. Launay, N. Buron, K. Begriche, A. Borgne-Sanchez, B. Fromenty, Drug-induced hepatic steatosis in absence of severe mitochondrial dysfunction in HepaRG cells: proof of multiple mechanism-based toxicity, Cell Biol Toxicol, 37 (2021) 151–175.

