## Supplementary Figures and Tables for "The Lipidome Landscape of Amiodarone Toxicity: An *in vivo* Lipid-centric Multi-Omics Study"

#: Corresponding to:

N.P.L.

### Supplementary Figures

**Supplementary Figure S1. Principal component analysis scores plots (with quality control group) of serum and liver lipidome of rats treated with amiodarone.** (A) Serum lipidome in ESI+ mode. (B) Serum lipidome in ESI- mode. (C) Liver lipidome in ESI+ mode. (D) Liver lipidome in ESI- mode. Abbreviations; ESI+, positive ion mode; ESI-, negative ion mode.

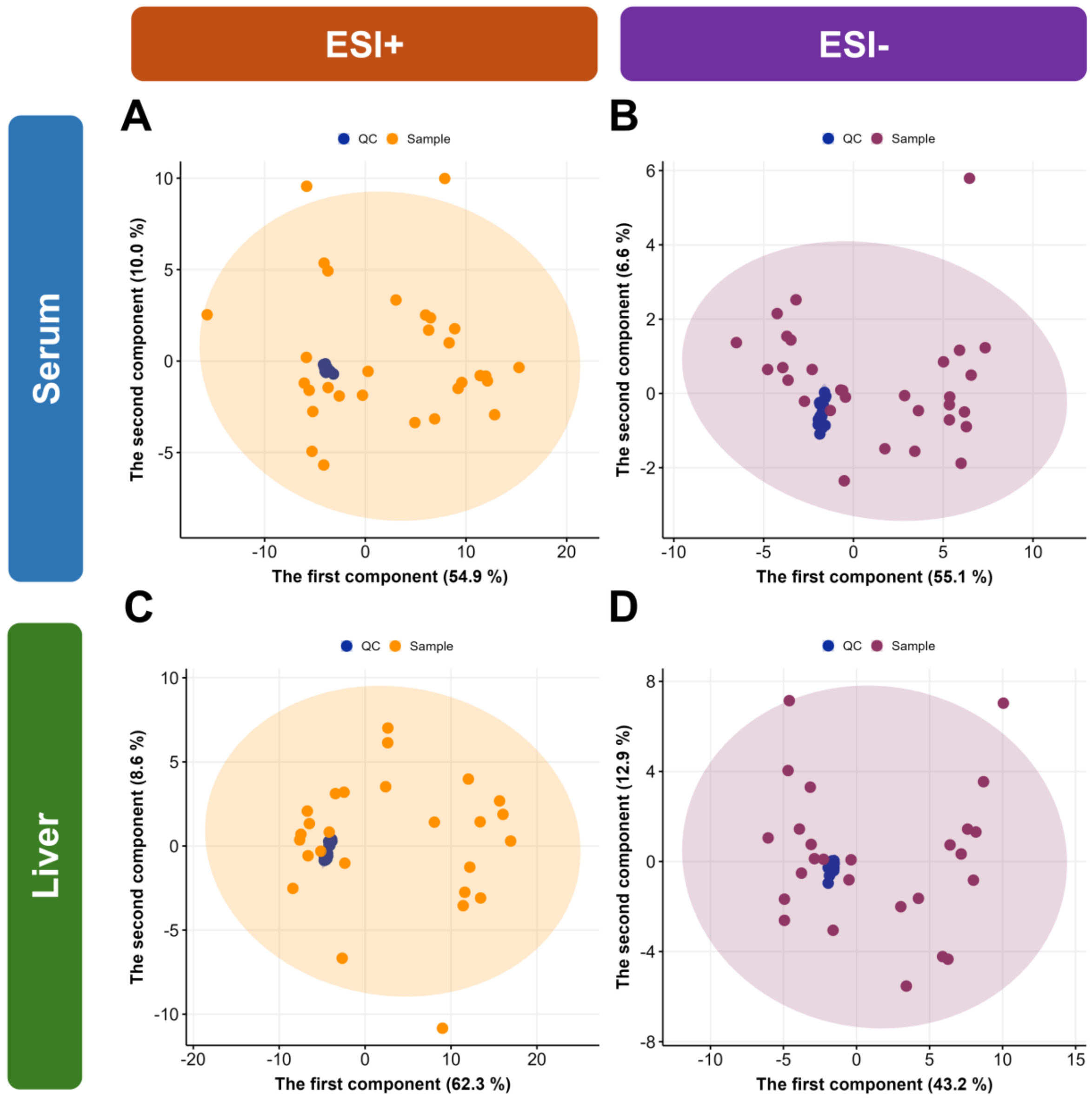

**Supplementary Figure S2. Principal component analysis scores plots of serum and liver lipidome of rats treated with amiodarone.** (A) Serum lipidome in ESI+ mode. (B) Serum lipidome in ESI- mode. (C) Liver lipidome in ESI+ mode. (D) Liver lipidome in ESI- mode. Abbreviations; ESI+, positive ion mode; ESI-, negative ion mode.

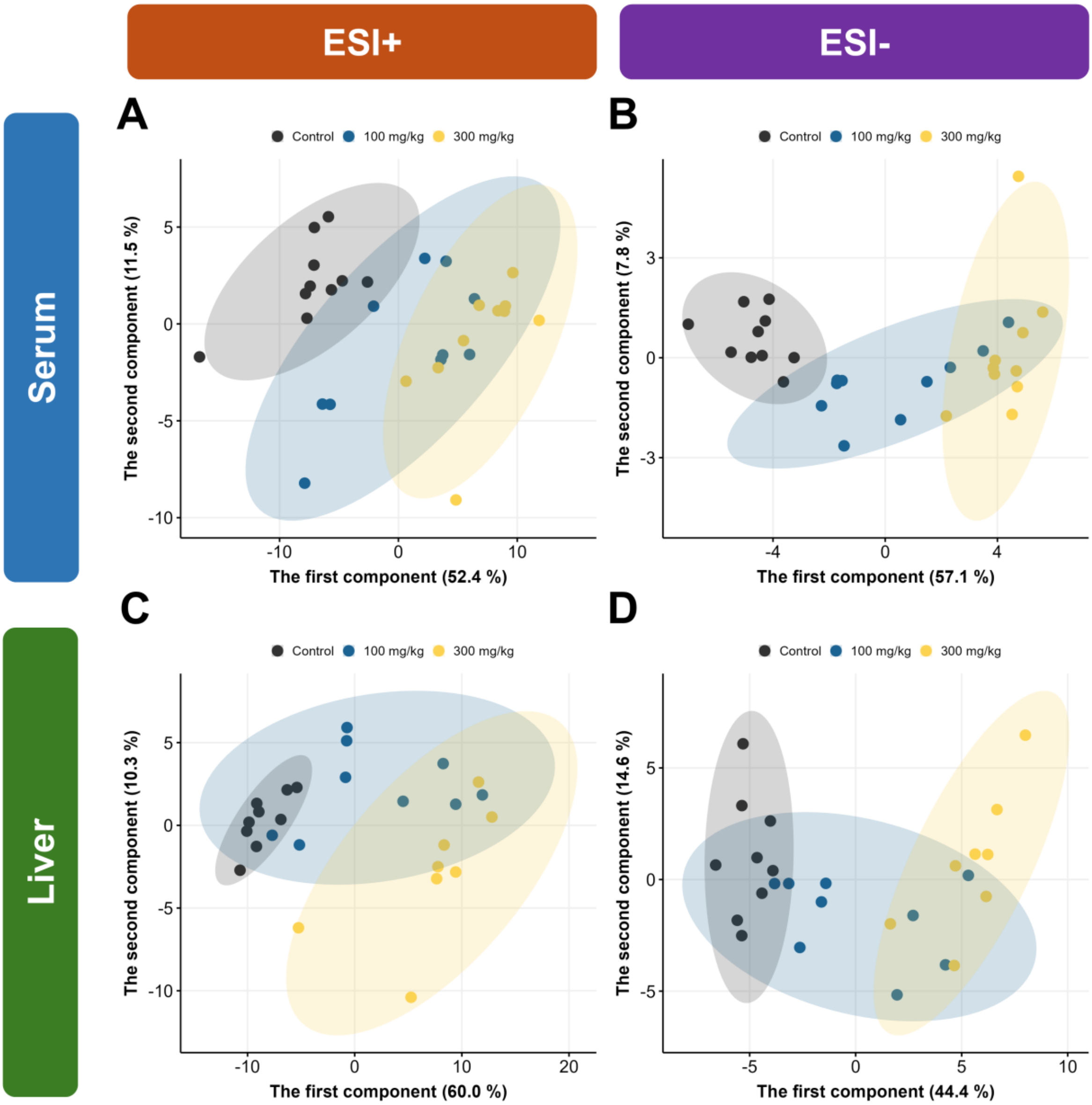

**Supplementary Figure S3. Partial least squares – discriminant analysis five-fold cross-validation results.** (A) Serum lipidome in ESI+ mode. (B) Serum lipidome in ESI- mode. (C) Liver lipidome in ESI+ mode. (D) Liver lipidome in ESI- mode. Abbreviations; ESI+, positive ion mode; ESI-, negative ion mode.

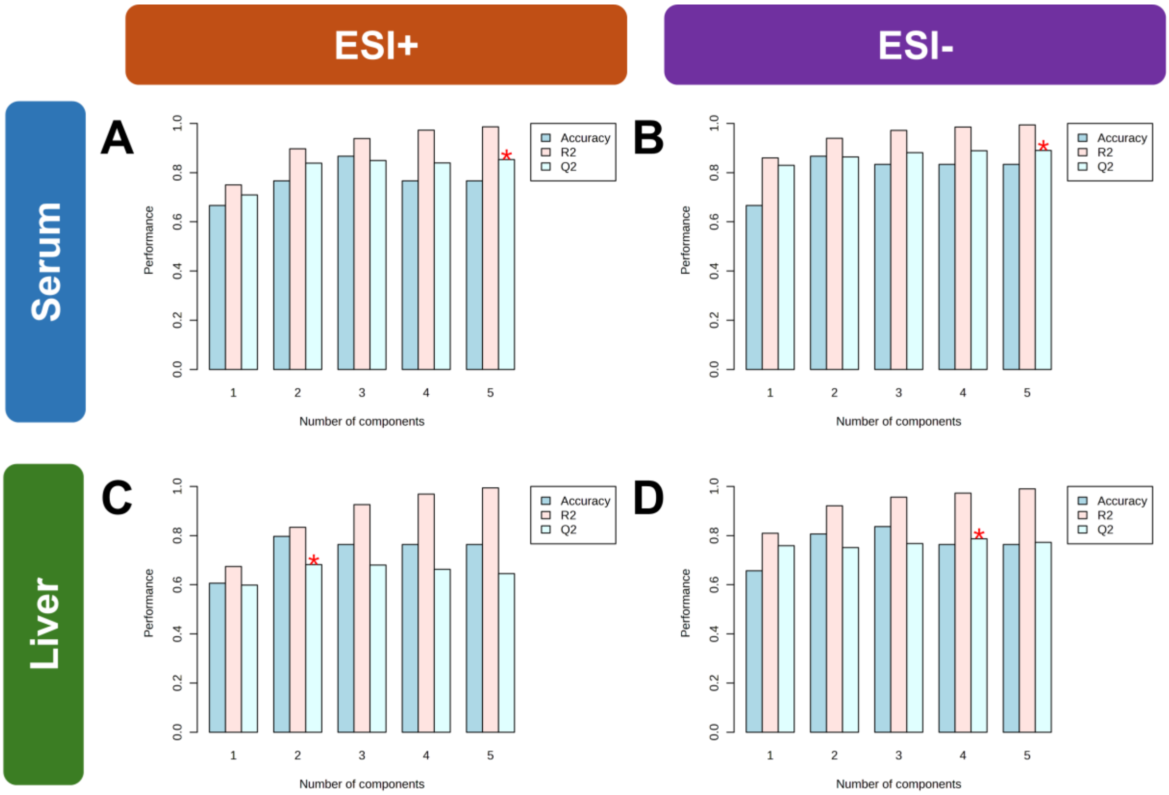

**Supplementary Figure S4. Volcano plots of differential lipids in the serum.** (A) ESI+ mode, 100 mg/kg vs. control. (B) ESI+ mode, 300 mg/kg vs. control. (C) ESI+ mode, 300 mg/kg vs. 100 mg/kg. (D) ESI- mode, 100 mg/kg vs. control. (E) ESI- mode, 300 mg/kg vs. control. (F) ESI- mode, 300 mg/kg vs. 100 mg/kg. Abbreviation: ESI+, positive ion mode; ESI-, negative ion mode; FC, fold change; FDR: false discovery rate.

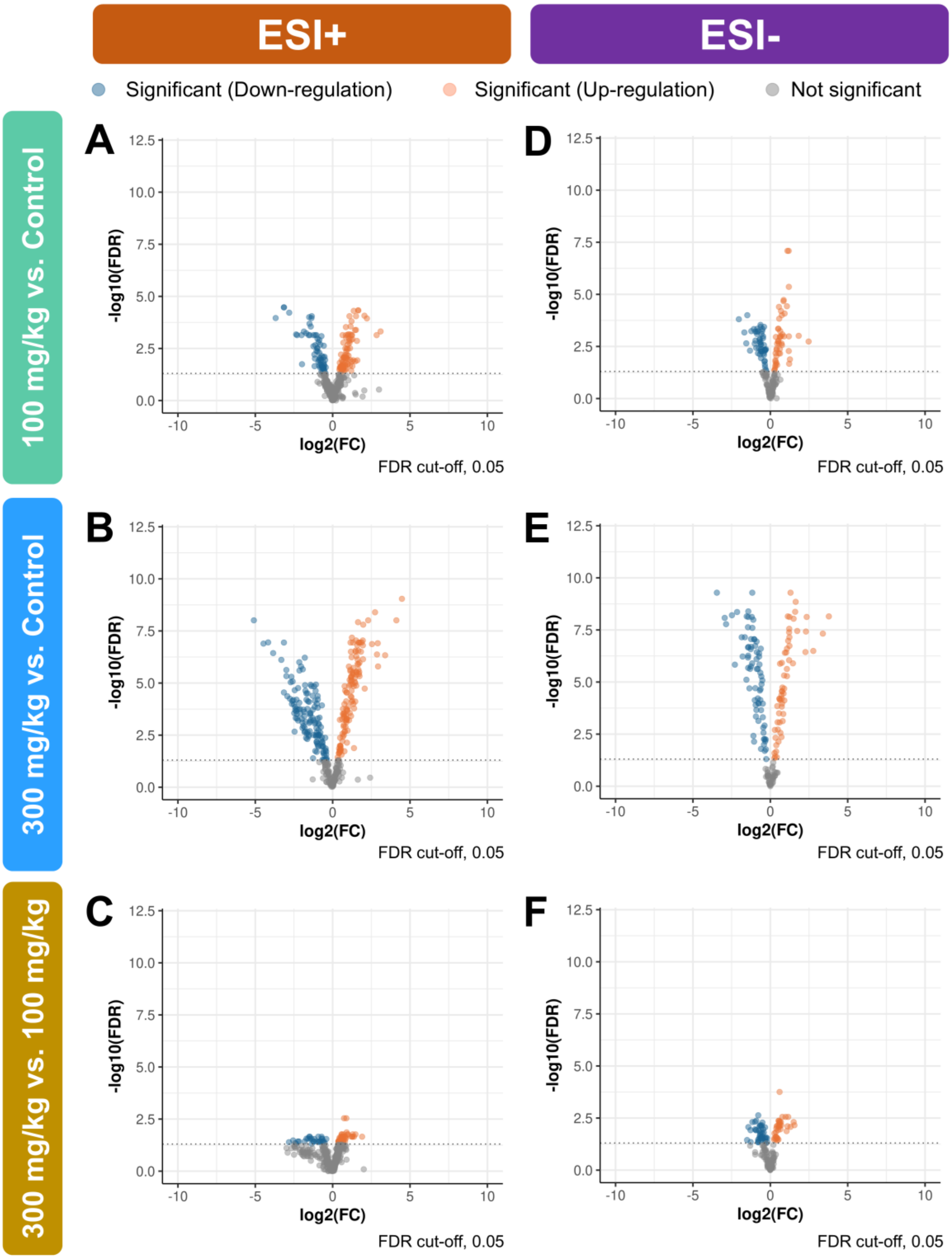

**Supplementary Figure S5. Volcano plots of differential lipids in the liver.** (A) ESI+ mode, 100 mg/kg vs. control. (B) ESI+ mode, 300 mg/kg vs. control. (C) ESI+ mode, 300 mg/kg vs. 100 mg/kg. (D) ESI- mode, 100 mg/kg vs. control. (E) ESI- mode, 300 mg/kg vs. control. (F) ESI- mode, 300 mg/kg vs. 100 mg/kg. Abbreviation: ESI+, positive ion mode; ESI-, negative ion mode; FC, fold change; FDR: false discovery rate.

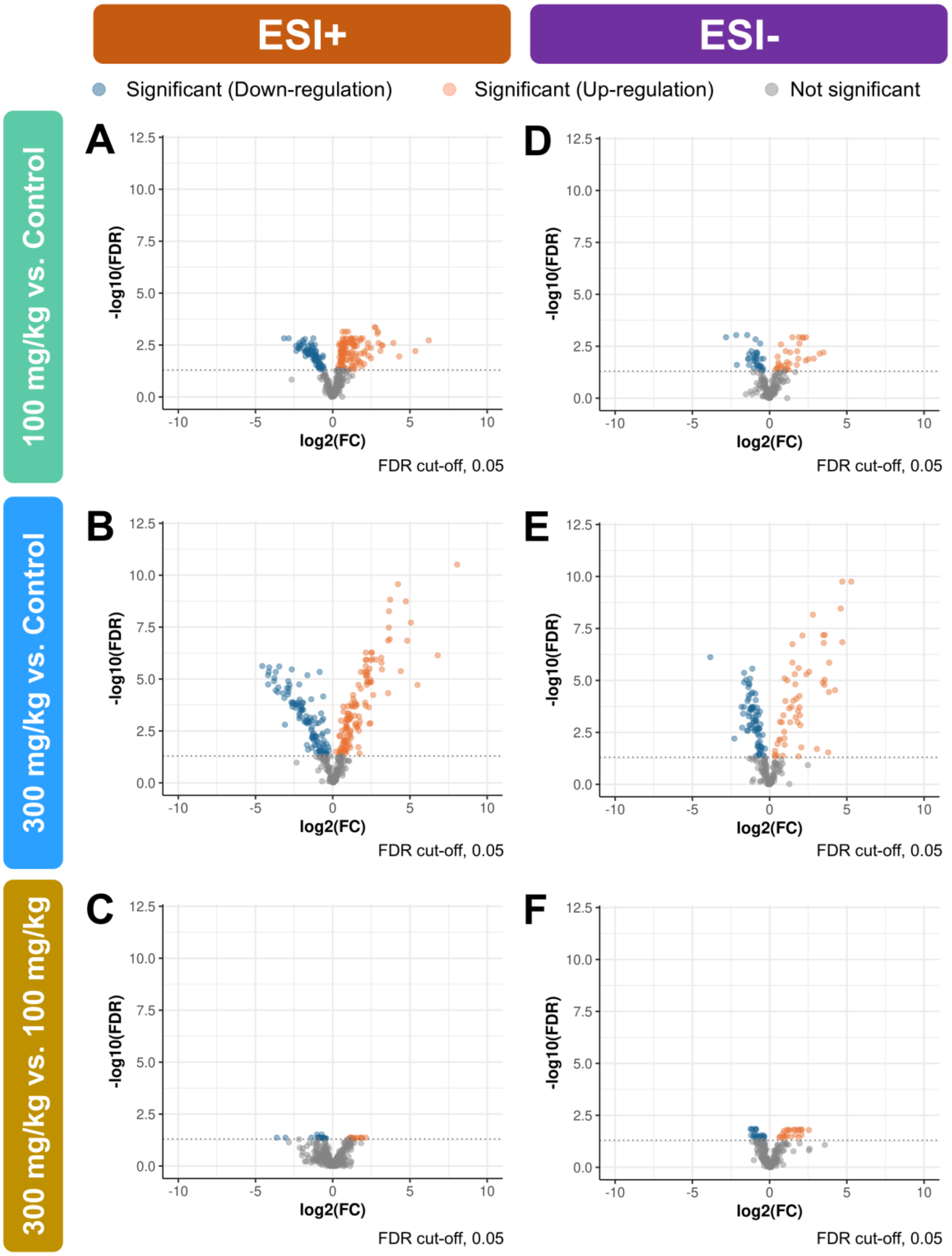

**Supplementary Figure S6. The disturbance of lipid metabolism and function under amiodarone exposure.** (A) The alteration of the abundance of lipid subclasses in the serum. (B) Lipid Ontology heatmap of systemic lipidome. (C) The alteration of the abundance of lipid subclasses in the liver. (D) Lipid Ontology heatmap of hepatic lipidome. Abbreviations; BMP, bismonoacylglycerophosphates; CAR, acylcarnitines; CE, cholesteryl esters; Cer, ceramides; CL, cardiolipins; DG, diacylglycerols; FA, fatty acids; LPC, lysophosphatidylcholines; PC, phosphatidylcholines; NAE, N-acyl ethanolamines; PC(O-), ether-linked phosphatidylcholines; PE, phosphatidylethanolamines; PE(O-), ether-linked phosphatidylethanolamines; PG, phosphatidylglycerols; PI, phosphatidylinositols; PS, phosphatidylserines; SM, sphingomyelins; TG, triacylglycerols; TG(O-), ether-linked triacylglycerols.

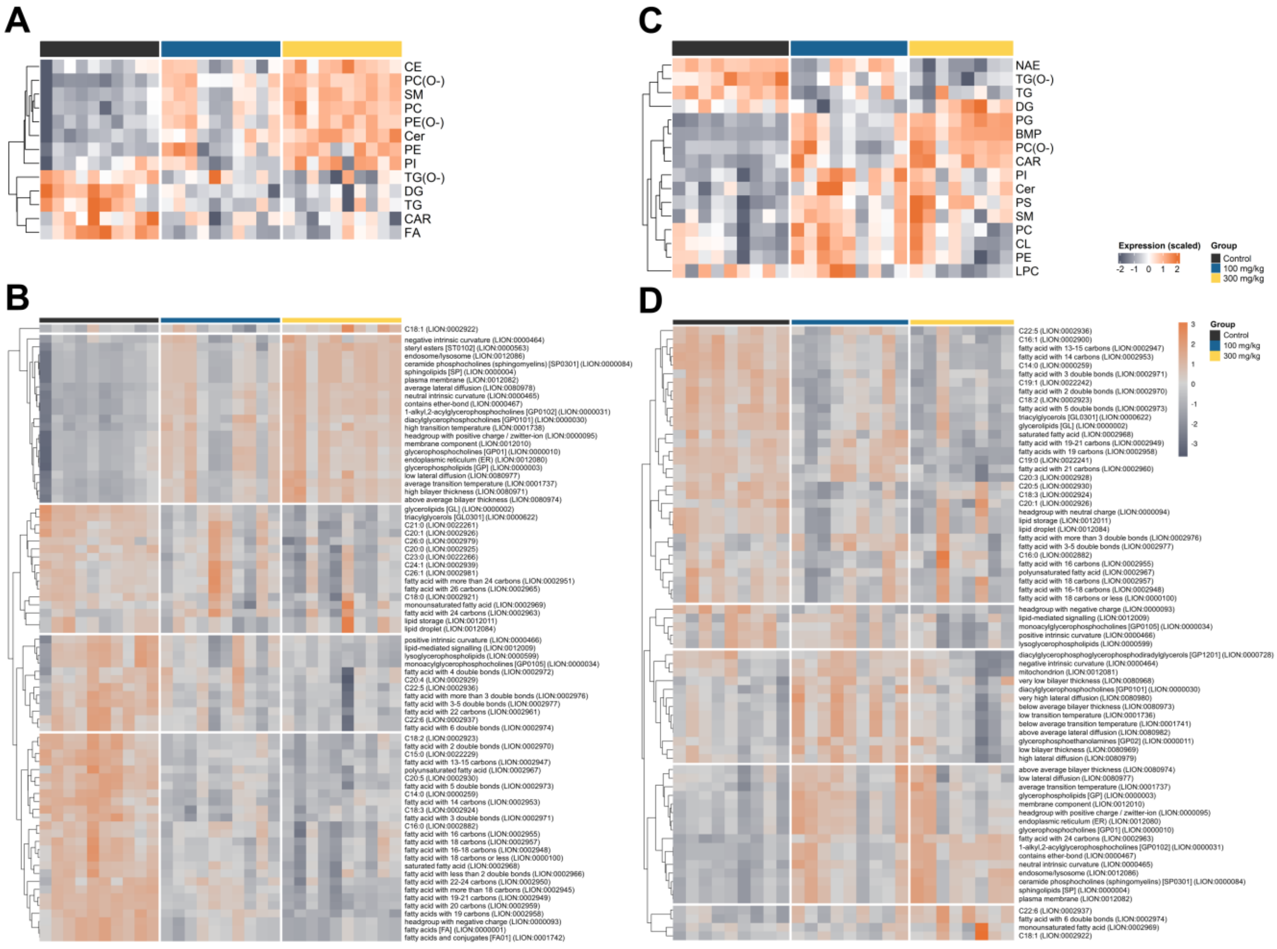

**Supplementary Figure S7. Systemic lipidome alterations in total acyl chain length and total double bonds of each lipid subclass, 100 mg/kg vs. control.** Abbreviations; CAR, acylcarnitines; CE, cholesteryl esters; Cer, ceramides; DG, diacylglycerols; FA, fatty acids; LPC, lysophosphatidylcholines; LPE, lysophosphatidylethanolamines; LPE(O-), ether-linked lysophosphatidylethanolamines; PC, phosphatidylcholines; PC(O-), ether-linked phosphatidylcholines; PE, phosphatidylethanolamines; PE(O-), ether-linked phosphatidylethanolamines; PI, phosphatidylinositols; SM, sphingomyelins; TG, triacylglycerols; TG(O-), ether-linked triacylglycerols; FC, fold change; FDR, false discovery rate; *, FDR < 0.05.

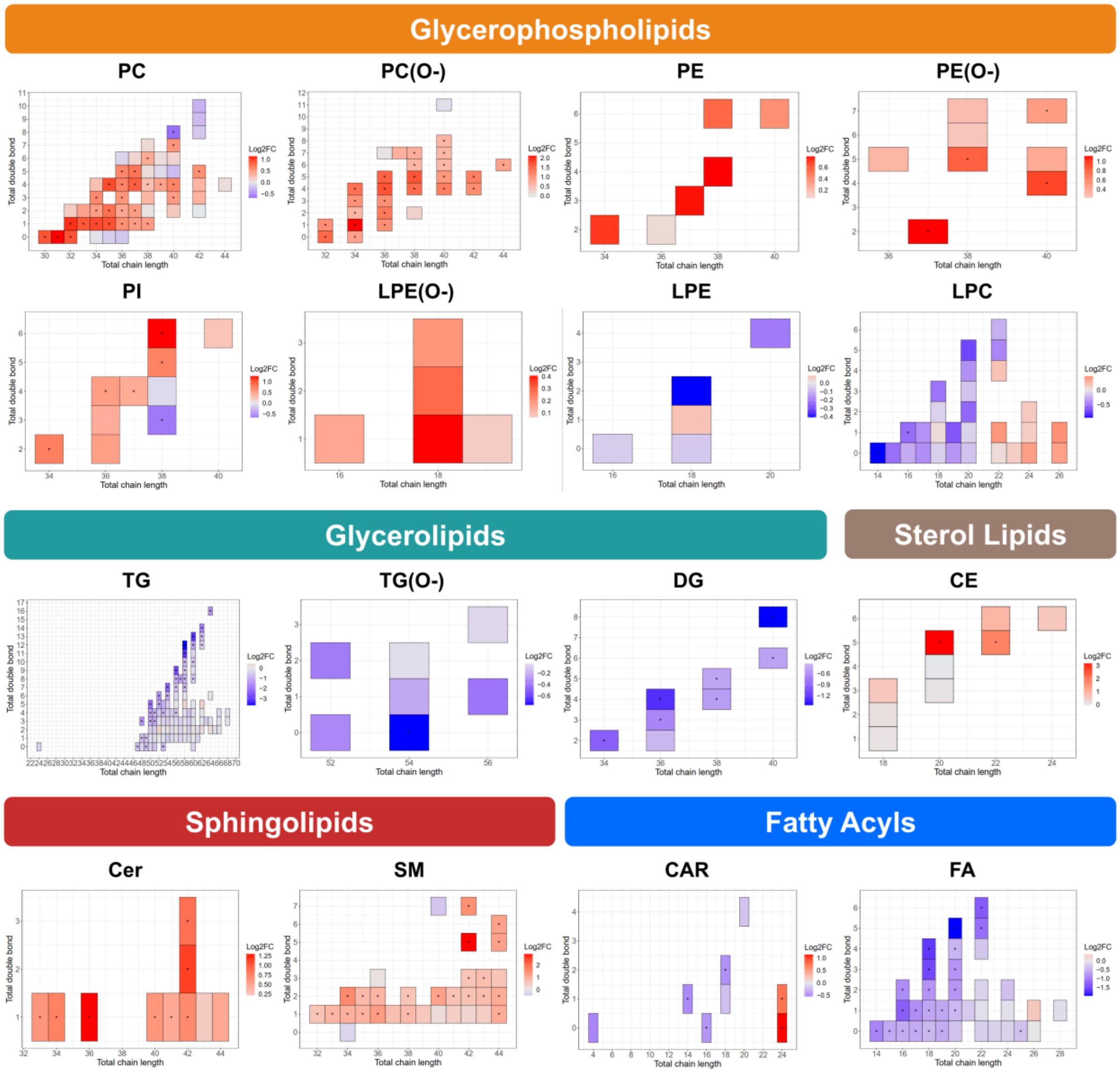

**Supplementary Figure S8. Systemic lipidome alterations in total acyl chain length and total double bonds of each lipid subclass, 300 mg/kg vs. control.** Abbreviations; CAR, acylcarnitine; CE, cholesteryl ester; Cer, ceramide; DG, diacylglycerol; FA, fatty acid; LPC, lysophosphatidylcholine; LPE, lysophosphatidylethanolamine; LPE(O-), ether-linked lysophosphatidylethanolamine; PC, phosphatidylcholine; PC(O-), ether-linked phosphatidylcholine; PE, phosphatidylethanolamine; PE(O-), ether-linked phosphatidylethanolamine; PI, phosphatidylinositol; SM, sphingomyelin; TG, triacylglycerol; TG(O-), ether-linked triacylglycerol; FC, fold change; FDR, false discovery rate; *, FDR < 0.05.

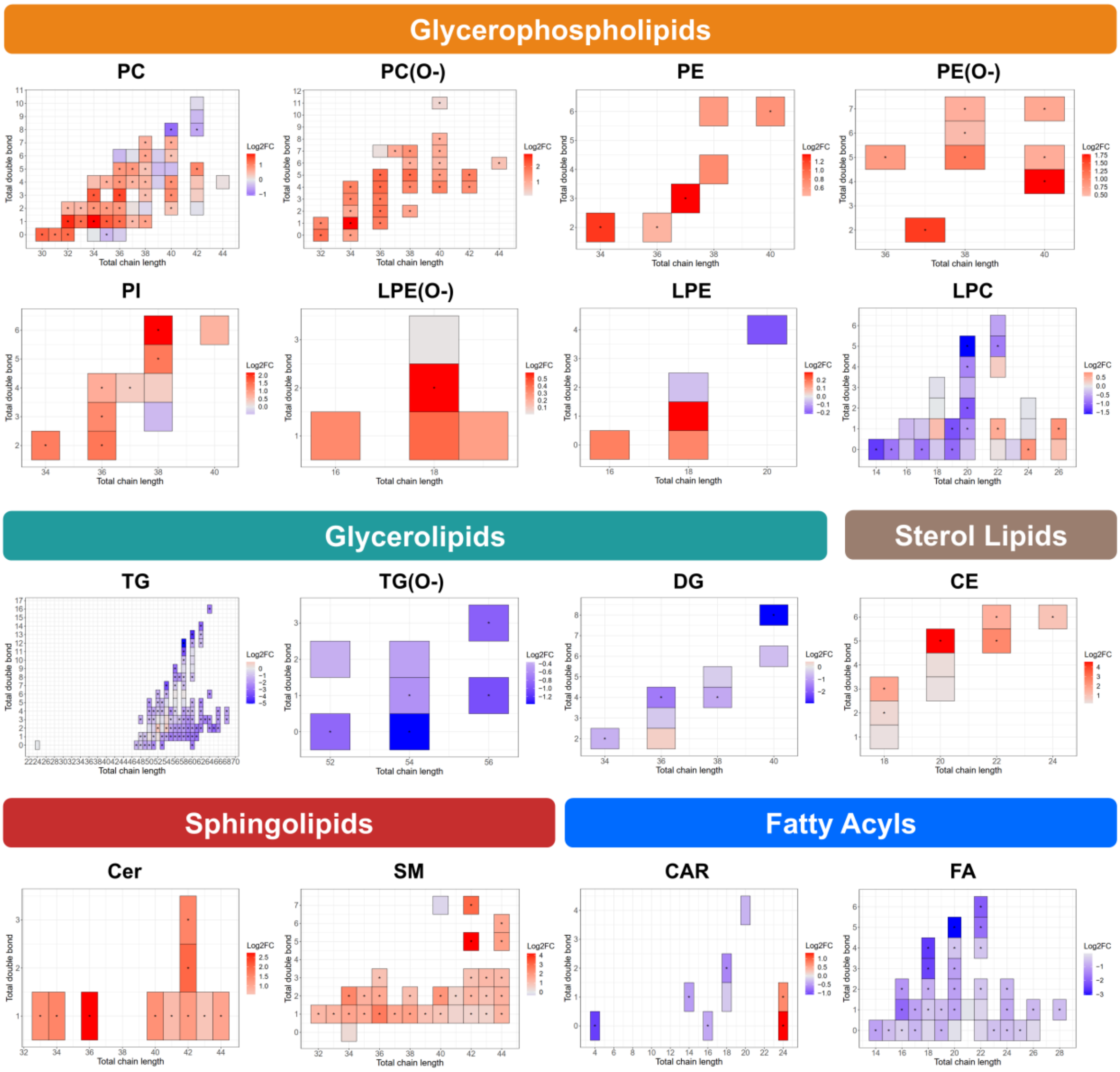

**Supplementary Figure S9. Hepatic lipidome alterations in total acyl chain length and total double bonds of each lipid subclass, 100 mg/kg vs. control.** Abbreviations; BMP, bismonoacylglycerophosphates; CAR, acylcarnitines; CE, cholesteryl esters; Cer, ceramides; CL, cardiolipins; DG, diacylglycerols; DG(O-), ether-linked diacylglycerols; FA, fatty acids; LPC, lysophosphatidylcholines; LPE, lysophosphatidylethanolamines; NAE, N-acyl ethanolamines; PC, phosphatidylcholines; PC(O-), ether-linked phosphatidylcholines; PE, phosphatidylethanolamines; PE(O-), ether-linked phosphatidylethanolamines; PG, phosphatidylglycerols; PI, phosphatidylinositols; PS, phosphatidylserines; SM, sphingomyelins; TG, triacylglycerols; TG(O-), ether-linked triacylglycerols; FC, fold change; FDR, false discovery rate; *, FDR < 0.05.

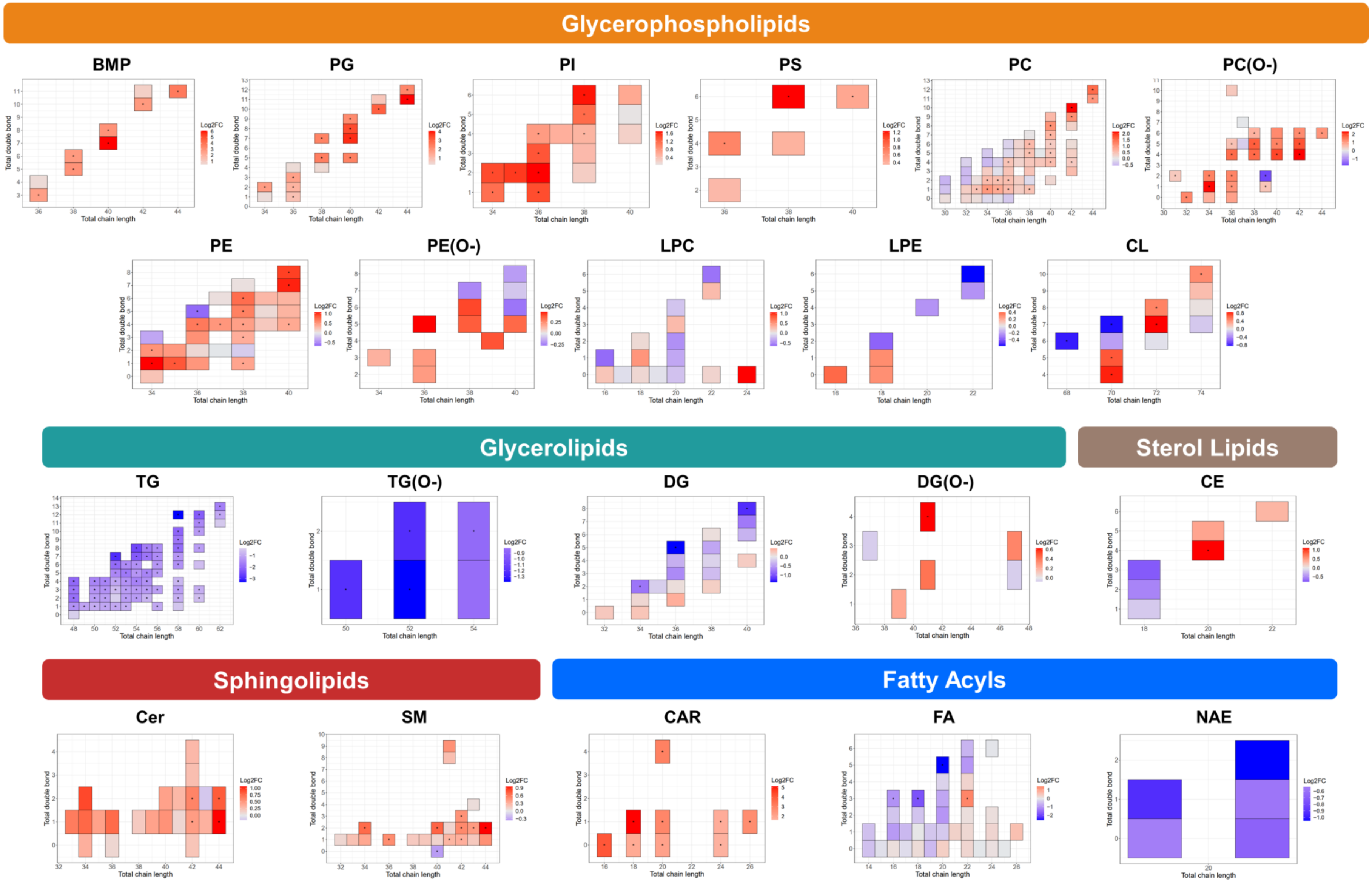

Glycerophospholipids
BMP
PG
PI
PS
PC
PC(O-)
PE
PE(O-)
LPC
LPE
CL
Glycerolipids
Sterol Lipids
TG
TG(O-)
DG
DG(O-)
CE
Sphingolipids
Fatty Acyls
Cer
SM
CAR
FA
NAE
Total chain length
Total double bond
Log2FC

**Supplementary Figure S10. Hepatic lipidome alterations in total acyl chain length and total double bonds of each lipid subclass, 300 mg/kg vs. control.** Abbreviations; BMP, bismonoacylglycerophosphates; CAR, acylcarnitines; CE, cholesteryl esters; Cer, ceramides; CL, cardiolipins; DG, diacylglycerols; DG(O-), ether-linked diacylglycerols; FA, fatty acid; LPC, lysophosphatidylcholines; LPE, lysophosphatidylethanolamines; NAE, N-acyl ethanolamines; PC, phosphatidylcholines; PC(O-), ether-linked phosphatidylcholines; PE, phosphatidylethanolamines; PE(O-), ether-linked phosphatidylethanolamines; PG, phosphatidylglycerols; PI, phosphatidylinositols; PS, phosphatidylserines; SM, sphingomyelins; TG, triacylglycerols; TG(O-), ether-linked triacylglycerols; FC, fold change; FDR, false discovery rate; *, FDR < 0.05.

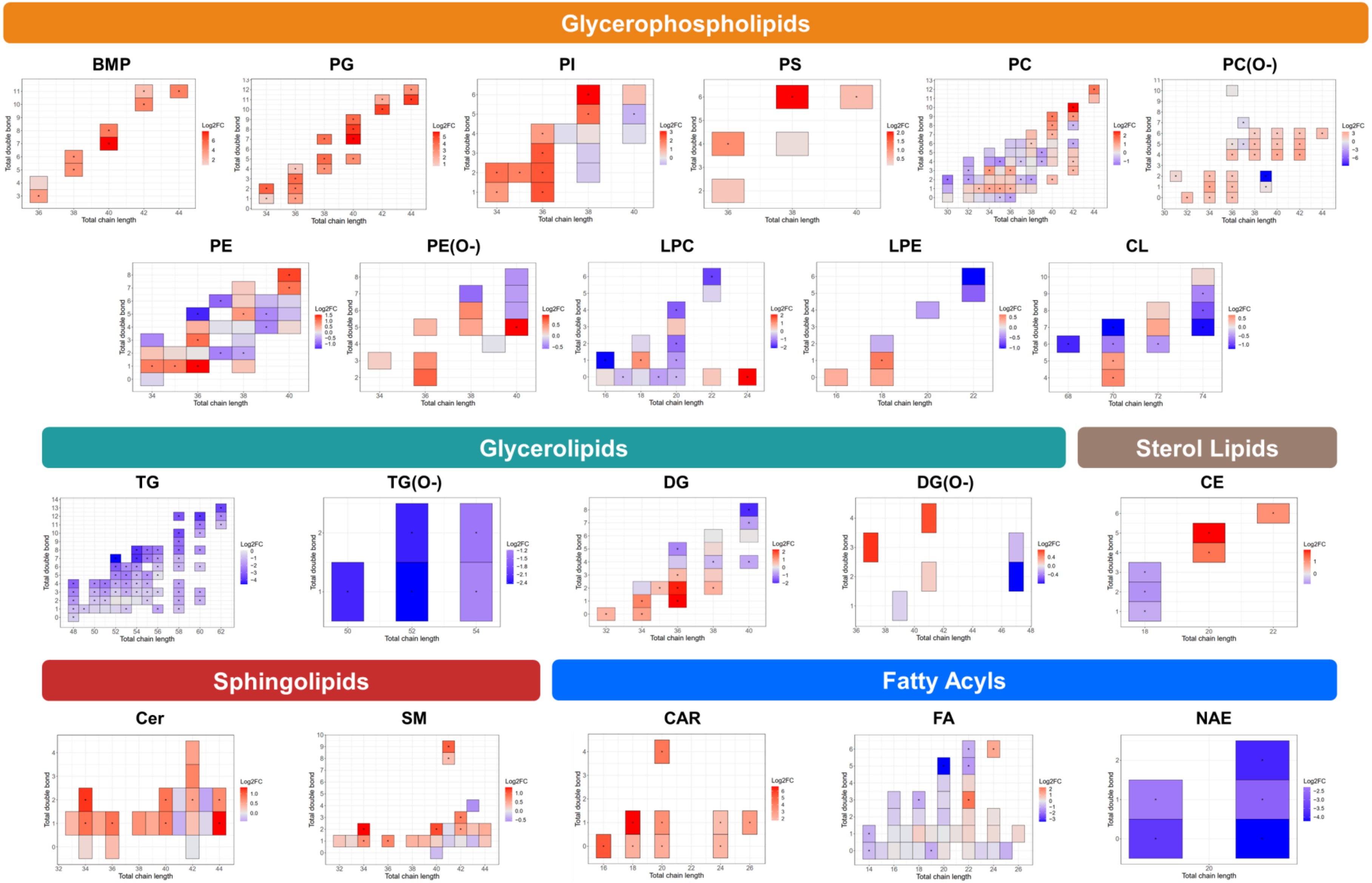
Glycerophospholipids
BMP
PG
PI
PS
PC
PC(O-)
PE
PE(O-)
LPC
LPE
CL
Glycerolipids
Sterol Lipids
TG
TG(O-)
DG
DG(O-)
CE
Sphingolipids
Fatty Acyls
Cer
SM
CAR
FA
NAE
Total chain length
Total double bond
Log2FC

### Supplementary Tables

**Supplementary Table S1. Number of differential lipids and lipid subclasses in two biospecimens of amiodarone-treated rats compared with the control group.**

| Comparison | | Number of differential lipid species | | Number of altered lipid subclasses* | | Altered lipid subclasses* (number of differential lipid species) | |
|---|---|---|---|---|---|---|---|
| | | Serum | Liver | Serum | Liver | Serum | Liver |
| 100 mg/kg vs. controls | ESI+ | 151 (85 up- and 66 down-regulation) | 223 (134 up and 89 down-regulation) | 7 | 10 | TG (49), PC(O-) (27), PC (24), SM (19), LPC (7), DG (5), CAR (5) | TG (77), PC (34), PC(O-) (24), PE (15), SM (12), CAR (9), BMP (6), Cer (6), PE(O-) (6), PI (6) |
| | ESI- | 99 (48 up and 51 down-regulation) | 67 (32 up and 35 down-regulation) | 7 | 4 | FA (28), LPC (12), SM (12), PC (11), PC(O-) (10), Cer (8), PI (6) | FA (20), PG (14), PC (13), PE(O-) (5) |
| 300 mg/kg vs. controls | ESI+ | 265 (130 up and 135 down-regulation) | 244 (129 up and 115 down-regulation) | 11 | 13 | TG (101), PC (40), PC(O-) (32), SM (29), LPC (20), CAR (6), | TG (73), PC (41), PC(O-) (25), PE (13), DG (13), LPC (12), CAR (9), SM (8), BMP (7), Cer (7), |

| | | | | | | | |
|---|---|---|---|---|---|---|---|
| | | | | | | DG (6), CE (5), Cer (5), PI (5), TG(O-) (5) | PE(O-) (5), NAE (5), TG(O-) (5) |
| | ESI- | 134 (59 up and 75 down-regulation) | 132 (55 up and 77 down-regulation) | 8 | 9 | FA (37), LPC (20), PC (17), SM (17), Cer (10), PC(O-) (10), PI (6), PE(O-) (5) | PC (24), FA (19), PG (19), PE (16), PI (13), PC(O-) (8), CL (7), SM (7), PE(O-) (5) |
| 300 mg/kg vs. 100 mg/kg | ESI+ | 76 (46 up and 30 down-regulation) | 37 (15 up and 22 down-regulation) | 5 | 2 | TG (21), SM (14), PC (12), LPC (7), PC(O-) (7) | PC (8), BMP (5) |
| | ESI- | 80 (35 up and 45 down-regulation) | 48 (22 up and 26 down-regulation) | 6 | 3 | FA (20), LPC (14), SM (12), PC (11), Cer (6), PI (5) | PG (13), PC (8), PE (8) |

*: Only lipid subclasses with ≥ 5 significant species are included. Abbreviations; BMP, bismonoacylglycerophosphates; CAR, acylcarnitines; CE, cholesteryl esters; Cer, ceramides; CL, cardiolipins; DG, diacylglycerols; ESI+, positive ion mode; ESI-, negative ion mode; FA, fatty acids; LPC, lysophosphatidylcholines; NAE, N-acyl ethanolamines; PC(O-), ether-linked phosphatidylcholines; PC, phosphatidylcholines; PE, phosphatidylethanolamine; PE(O-), ether-linked phosphatidylethanolamines; PG, phosphatidylglycerols; PI, phosphatidylinositols; SM, sphingomyelins; TG(O-), ether-linked triacylglycerols; TG, triacylglycerols.

**Supplementary Table S2. Differential lipids in serum of amiodarone-treated rats compared with the control group.**

| No | Differential lipids | 100 mg/kg vs. control | | 300 mg/kg vs. control | |
|---|---|---|---|---|---|
| | | FC | FDR | FC | FDR |
| 1 | CAR(14:1) | 0.706 | 4.67E-02 | 0.716 | 4.00E-02 |
| 2 | CAR(16:0) | 0.737 | 1.21E-02 | 0.754 | 6.82E-03 |
| 3 | CAR(18:2) | 0.720 | 9.07E-03 | 0.648 | 1.38E-04 |
| 4 | CAR(24:0) | 2.195 | 1.07E-03 | 2.494 | 4.05E-08 |
| 5 | CAR(24:1) | 1.679 | 1.90E-02 | 1.648 | 1.13E-03 |
| 6 | CAR(4:0) | - | - | 0.477 | 1.17E-03 |
| 7 | CE(18:2) | - | - | 1.332 | 3.84E-02 |
| 8 | CE(18:3) | - | - | 3.479 | 2.32E-06 |
| 9 | CE(20:5) | 8.858 | 3.80E-04 | 23.460 | 3.81E-10 |
| 10 | CE(22:5)_1 | 4.831 | 9.78E-05 | 7.885 | 6.20E-08 |
| 11 | CE(22:6)_2 | 1.901 | 5.00E-02 | 2.889 | 4.71E-05 |
| 12 | CE(24:6) | - | - | 1.992 | 4.37E-03 |
| 13 | Cer(33:1;2O) | 1.541 | 2.01E-02 | 2.678 | 1.66E-06 |
| 14 | Cer(34:1;2O)\|Cer(18:1;2O/16:0) | 1.700 | 3.21E-03 | 2.957 | 7.48E-07 |
| 15 | Cer(36:1;2O)\|Cer(18:1;2O/18:0) | 2.452 | 2.87E-03 | 6.663 | 7.91E-08 |
| 16 | Cer(40:1;2O) | - | - | 1.674 | 1.38E-04 |
| 17 | Cer(40:1;2O)\|Cer(18:1;2O/22:0) | 1.607 | 1.28E-02 | 3.011 | 1.19E-06 |
| 18 | Cer(41:1;2O)\|Cer(18:1;2O/23:0) | - | - | 1.656 | 3.96E-04 |
| 19 | Cer(42:1;2O)\|Cer(18:1;2O/24:0) | 1.628 | 1.39E-02 | 2.365 | 1.07E-05 |
| 20 | Cer(42:2;2O)\|Cer(18:1;2O/24:1) | 2.209 | 6.83E-04 | 3.942 | 8.54E-08 |
| 21 | Cer(42:3;2O)\|Cer(18:1;2O/24:2) | 1.842 | 6.83E-03 | 2.995 | 7.48E-07 |
| 22 | Cer(43:1;2O)\|Cer(18:1;2O/25:0) | - | - | 1.481 | 2.26E-02 |
| 23 | Cer(44:1;2O)\|Cer(18:1;2O/26:0) | - | - | 2.130 | 1.92E-04 |
| 24 | DG(34:2) | 0.516 | 3.09E-03 | 0.589 | 7.06E-03 |
| 25 | DG(36:4)\|DG(18:2_18:2) | 0.351 | 8.47E-04 | 0.288 | 3.04E-04 |
| 26 | DG(38:4)\|DG(18:0_20:4) | 0.694 | 1.34E-02 | 0.539 | 1.53E-04 |
| 27 | DG(40:8) | 0.366 | 6.83E-04 | 0.132 | 6.77E-06 |
| 28 | FA(14:0) | 0.521 | 8.47E-04 | 0.495 | 5.26E-05 |
| 29 | FA(15:0)_2 | 0.610 | 9.97E-04 | 0.572 | 5.26E-05 |
| 30 | FA(16:0) | 0.683 | 2.76E-03 | 0.788 | 3.36E-02 |
| 31 | FA(16:1) | 0.411 | 2.93E-03 | 0.270 | 2.89E-06 |

| | | | | | |
|---|---|---|---|---|---|
| 32 | FA(16:2) | 0.552 | 3.97E-04 | 0.430 | 2.20E-07 |
| 33 | FA(17:0)_1 | 0.630 | 2.17E-03 | 0.493 | 1.98E-06 |
| 34 | FA(17:0)_2 | 0.687 | 3.82E-03 | 0.637 | 4.31E-04 |
| 35 | FA(17:1) | 0.559 | 9.05E-04 | 0.494 | 2.10E-06 |
| 36 | FA(18:1) | 0.657 | 6.62E-04 | 0.723 | 2.73E-03 |
| 37 | FA(18:2) | 0.582 | 3.93E-04 | 0.557 | 1.15E-05 |
| 38 | FA(18:3) | 0.356 | 8.47E-04 | 0.189 | 1.14E-08 |
| 39 | FA(18:4) | 0.323 | 3.12E-04 | 0.173 | 2.99E-08 |
| 40 | FA(19:0) | 0.681 | 7.41E-03 | 0.578 | 1.05E-04 |
| 41 | FA(19:1) | 0.655 | 6.62E-04 | 0.645 | 2.80E-04 |
| 42 | FA(20:1) | 0.738 | 1.08E-02 | 0.509 | 1.98E-06 |
| 43 | FA(20:2) | 0.589 | 5.81E-04 | 0.489 | 1.19E-06 |
| 44 | FA(20:3) | 0.583 | 8.47E-04 | 0.484 | 3.69E-06 |
| 45 | FA(20:4) | 0.718 | 1.34E-02 | 0.701 | 1.52E-03 |
| 46 | FA(20:5) | 0.258 | 6.17E-05 | 0.122 | 3.81E-10 |
| 47 | FA(22:2) | - | - | 0.701 | 7.23E-03 |
| 48 | FA(22:4) | - | - | 0.701 | 1.95E-03 |
| 49 | FA(22:5) | 0.452 | 7.08E-04 | 0.291 | 6.86E-08 |
| 50 | FA(22:6) | 0.395 | 4.27E-05 | 0.242 | 2.60E-09 |
| 51 | FA(23:0) | - | - | 0.672 | 2.15E-03 |
| 52 | FA(24:0) | - | - | 0.657 | 1.77E-03 |
| 53 | FA(24:1) | - | - | 0.525 | 2.38E-04 |
| 54 | FA(24:2) | - | - | 0.593 | 2.11E-04 |
| 55 | FA(25:0) | - | - | 0.731 | 2.09E-02 |
| 56 | FA(26:1) | - | - | 0.624 | 2.13E-02 |
| 57 | FA(28:1) | - | - | 0.653 | 3.08E-02 |
| 58 | LPC(14:0-SN2) | 0.523 | 8.72E-04 | 0.403 | 5.98E-06 |
| 59 | LPC(15:0-SN2) | 0.686 | 3.06E-02 | 0.541 | 5.26E-05 |
| 60 | LPC(16:1-SN1) | 0.682 | 7.71E-03 | 0.788 | 3.44E-02 |
| 61 | LPC(17:0-SN1) | - | - | 0.537 | 3.33E-05 |
| 62 | LPC(18:0-SN1)_1 | - | - | 0.770 | 2.56E-02 |
| 63 | LPC(18:3-SN2) | 0.695 | 2.79E-02 | - | - |
| 64 | LPC(19:0) | - | - | 0.565 | 2.09E-04 |
| 65 | LPC(19:0-SN1) | - | - | 0.492 | 3.24E-05 |
| 66 | LPC(19:1) | 0.638 | 2.18E-02 | 0.448 | 9.16E-04 |
| 67 | LPC(20:1-SN1)_1 | - | - | 0.532 | 3.50E-04 |

| | | | | | |
|---|---|---|---|---|---|
| 68 | LPC(20:1-SN1)_2 | - | - | 0.645 | 5.30E-03 |
| 69 | LPC(20:2-SN1) | 0.667 | 3.18E-02 | 0.458 | 1.99E-05 |
| 70 | LPC(20:4-SN1) | - | - | 0.587 | 4.84E-04 |
| 71 | LPC(20:5-SN1)_2 | 0.475 | 6.04E-04 | 0.303 | 4.87E-07 |
| 72 | LPC(22:1-SN1) | - | - | 1.490 | 3.41E-03 |
| 73 | LPC(22:5-SN1) | - | - | 0.566 | 4.69E-03 |
| 74 | LPC(22:6-SN1) | - | - | 0.696 | 3.59E-02 |
| 75 | LPC(24:0-SN1) | - | - | 1.732 | 1.94E-04 |
| 76 | LPC(26:1-SN2) | - | - | 1.724 | 1.40E-03 |
| 77 | LPE(18:1) | - | - | 1.229 | 4.06E-02 |
| 78 | LPE(18:2) | 0.756 | 4.12E-02 | - | - |
| 79 | LPE(O-18:2) | - | - | 1.498 | 3.13E-03 |
| 80 | PC(30:0) | 1.882 | 5.19E-04 | 2.461 | 1.39E-07 |
| 81 | PC(31:0) | 2.194 | 5.45E-05 | 2.166 | 2.85E-07 |
| 82 | PC(32:0)\|PC(16:0_16:0) | 1.886 | 5.95E-04 | 2.384 | 8.54E-08 |
| 83 | PC(32:1)\|PC(16:0_16:1) | 2.073 | 3.97E-04 | 3.171 | 1.09E-07 |
| 84 | PC(32:2)\|PC(14:0_18:2) | - | - | 1.453 | 5.31E-03 |
| 85 | PC(33:1) | 1.864 | 1.75E-04 | 2.339 | 6.86E-08 |
| 86 | PC(33:2) | - | - | 1.400 | 4.21E-03 |
| 87 | PC(34:1)\|PC(16:0_18:1) | 1.952 | 5.75E-03 | 3.382 | 4.78E-08 |
| 88 | PC(34:2)\|PC(16:0_18:2) | 1.409 | 2.07E-02 | 1.850 | 8.12E-06 |
| 89 | PC(34:3)\|PC(16:1_18:2) | 1.539 | 9.10E-03 | 2.610 | 1.18E-07 |
| 90 | PC(35:0) | - | - | 0.762 | 3.69E-02 |
| 91 | PC(35:1)\|PC(17:0_18:1) | 1.896 | 2.87E-03 | 2.607 | 1.76E-06 |
| 92 | PC(35:2)\|PC(17:0_18:2) | 1.704 | 6.38E-03 | 1.750 | 1.54E-04 |
| 93 | PC(35:4) | 1.936 | 2.02E-03 | 1.502 | 1.12E-02 |
| 94 | PC(36:1)\|PC(18:0_18:1) | 1.503 | 2.78E-02 | 2.330 | 1.35E-05 |
| 95 | PC(36:2)\|PC(18:0_18:2) | 1.351 | 3.68E-02 | 1.776 | 3.70E-05 |
| 96 | PC(36:3)\|PC(18:1_18:2) | 1.456 | 1.83E-02 | 2.855 | 3.55E-06 |
| 97 | PC(36:4)\|PC(16:0_20:4) | 1.606 | 3.61E-04 | 1.801 | 1.07E-05 |
| 98 | PC(36:4)\|PC(18:2_18:2) | - | - | 2.001 | 7.48E-05 |
| 99 | PC(36:5)_2 | - | - | 1.930 | 1.19E-04 |
| 100 | PC(36:5)\|PC(16:1_20:4) | 1.823 | 2.84E-02 | 1.818 | 1.35E-03 |
| 101 | PC(37:1) | 1.417 | 1.81E-02 | 1.406 | 1.91E-02 |
| 102 | PC(37:2)\|PC(19:0_18:2) | 1.424 | 5.00E-02 | - | - |
| 103 | PC(37:4)\|PC(17:0_20:4) | 1.815 | 2.03E-03 | 1.464 | 2.61E-03 |

| | | | | | |
|---|---|---|---|---|---|
| 104 | PC(37:5) | 1.666 | 8.80E-04 | 1.769 | 1.61E-04 |
| 105 | PC(38:1) | 1.423 | 1.34E-02 | 1.648 | 9.50E-05 |
| 106 | PC(38:3)\|PC(18:0_20:3) | - | - | 1.351 | 2.67E-02 |
| 107 | PC(38:4) | 1.683 | 4.62E-03 | 2.407 | 1.76E-06 |
| 108 | PC(38:4)\|PC(18:0_20:4) | - | - | 1.448 | 1.74E-04 |
| 109 | PC(38:5) | - | - | 1.471 | 3.49E-03 |
| 110 | PC(38:5)\|PC(16:0_22:5) | - | - | 1.458 | 1.19E-02 |
| 111 | PC(38:5)\|PC(18:1_20:4) | - | - | 1.308 | 2.36E-02 |
| 112 | PC(38:6) | 1.351 | 3.84E-02 | 1.674 | 1.48E-04 |
| 113 | PC(38:7) | - | - | 1.496 | 1.33E-02 |
| 114 | PC(39:4) | 1.373 | 3.69E-02 | - | - |
| 115 | PC(40:2) | - | - | 1.318 | 1.33E-02 |
| 116 | PC(40:3) | 1.425 | 8.63E-03 | 1.940 | 1.10E-05 |
| 117 | PC(40:4)\|PC(18:0_22:4) | 1.578 | 2.00E-02 | 1.877 | 7.69E-04 |
| 118 | PC(40:6) | - | - | 0.431 | 9.01E-05 |
| 119 | PC(40:6)\|PC(18:0_22:6) | - | - | 1.332 | 2.65E-02 |
| 120 | PC(40:7)\|PC(18:1_22:6) | - | - | 1.604 | 2.84E-04 |
| 121 | PC(40:8)\|PC(20:4_20:4) | 0.632 | 5.41E-03 | 0.481 | 1.74E-04 |
| 122 | PC(42:5) | 1.499 | 2.87E-03 | 1.985 | 1.68E-05 |
| 123 | PC(42:8) | - | - | 0.679 | 2.71E-02 |
| 124 | PC(O-32:0)\|PC(O-16:0_16:0) | 2.817 | 2.97E-04 | 3.890 | 4.78E-08 |
| 125 | PC(O-32:1) | 2.310 | 1.72E-04 | 2.941 | 1.14E-07 |
| 126 | PC(O-32:1)\|PC(O-16:1_16:0) | 2.059 | 5.13E-04 | 3.004 | 3.10E-08 |
| 127 | PC(O-34:0) | 1.824 | 3.21E-03 | 2.344 | 3.55E-06 |
| 128 | PC(O-34:1)\|PC(O-18:1_16:0) | 4.305 | 5.45E-05 | 7.028 | 1.57E-09 |
| 129 | PC(O-34:2)\|PC(O-18:2_16:0) | 1.542 | 3.17E-02 | 2.118 | 1.27E-04 |
| 130 | PC(O-34:3) | 2.055 | 1.14E-03 | 3.081 | 1.42E-06 |
| 131 | PC(O-34:4) | 2.665 | 2.96E-05 | 3.207 | 2.72E-07 |
| 132 | PC(O-36:1) | 2.121 | 3.73E-03 | 3.355 | 9.34E-07 |
| 133 | PC(O-36:2) | 2.517 | 7.08E-04 | 3.740 | 1.14E-07 |
| 134 | PC(O-36:3) | - | - | 3.500 | 3.42E-07 |
| 135 | PC(O-36:4)\|PC(O-16:0_20:4) | 3.253 | 2.96E-05 | 4.090 | 1.14E-07 |
| 136 | PC(O-36:5)\|PC(O-16:1_20:4) | 2.946 | 5.45E-05 | 4.146 | 9.15E-09 |
| 137 | PC(O-37:7) | - | - | 1.976 | 2.89E-04 |
| 138 | PC(O-38:2) | - | - | 2.199 | 3.46E-04 |
| 139 | PC(O-38:4) | 3.103 | 1.03E-03 | 2.458 | 1.20E-05 |

| | | | | | |
|---|---|---|---|---|---|
| 140 | PC(O-38:4)\|PC(O-18:0_20:4) | 2.578 | 7.53E-03 | 3.366 | 7.89E-07 |
| 141 | PC(O-38:5)\|PC(O-18:1_20:4)_1 | 3.249 | 2.96E-05 | 4.076 | 4.73E-08 |
| 142 | PC(O-38:5)\|PC(O-18:1_20:4)_2 | 2.275 | 3.61E-04 | 2.414 | 2.01E-06 |
| 143 | PC(O-38:6) | 2.414 | 9.85E-05 | 2.666 | 6.84E-07 |
| 144 | PC(O-38:6)\|PC(O-18:2_20:4) | - | - | 1.734 | 6.48E-04 |
| 145 | PC(O-38:7) | - | - | 2.730 | 8.72E-05 |
| 146 | PC(O-40:4)_1 | 2.366 | 8.47E-04 | 2.447 | 2.24E-05 |
| 147 | PC(O-40:4)_2 | 2.471 | 4.48E-04 | 3.650 | 4.78E-08 |
| 148 | PC(O-40:5)_1 | 1.990 | 5.77E-03 | 2.696 | 7.01E-05 |
| 149 | PC(O-40:5)_2 | 1.957 | 3.48E-03 | 2.148 | 6.66E-05 |
| 150 | PC(O-40:6) | 1.517 | 1.20E-02 | 1.731 | 3.58E-04 |
| 151 | PC(O-40:7) | 1.537 | 1.90E-02 | 2.031 | 1.53E-04 |
| 152 | PC(O-40:8) | 1.548 | 1.78E-02 | 1.934 | 2.55E-05 |
| 153 | PC(O-42:4) | 2.156 | 6.62E-04 | 2.848 | 4.01E-06 |
| 154 | PC(O-42:5) | 2.402 | 5.19E-04 | 3.341 | 1.17E-07 |
| 155 | PC(O-44:6) | 1.794 | 1.29E-02 | 1.570 | 1.51E-02 |
| 156 | PE(34:2)\|PE(16:0_18:2) | - | - | 2.368 | 2.43E-04 |
| 157 | PE(36:2)\|PE(18:0_18:2) | - | - | 1.446 | 2.95E-02 |
| 158 | PE(36:2)\|PE(18:1_18:1) | - | - | 1.314 | 1.89E-02 |
| 159 | PE(37:3) | 1.716 | 2.52E-02 | 2.575 | 3.80E-06 |
| 160 | PE(40:6)\|PE(18:0_22:6)_2 | - | - | 2.646 | 7.68E-03 |
| 161 | PE(O-36:5)\|PE(O-16:1_20:4) | - | - | 1.744 | 1.16E-03 |
| 162 | PE(O-37:2) | 2.163 | 8.47E-04 | 3.021 | 2.02E-07 |
| 163 | PE(O-38:5)\|PE(O-18:1_20:4) | 1.782 | 1.34E-02 | 2.191 | 5.58E-06 |
| 164 | PE(O-38:6)\|PE(O-18:2_20:4) | - | - | 1.388 | 2.55E-02 |
| 165 | PE(O-38:7)\|PE(O-16:1_22:6) | - | - | 1.512 | 7.05E-03 |
| 166 | PE(O-40:4)\|PE(O-18:0_22:4) | 1.943 | 7.53E-03 | 3.430 | 4.21E-06 |
| 167 | PE(O-40:5)\|PE(O-18:1_22:4) | - | - | 1.535 | 1.20E-02 |
| 168 | PE(O-40:7)\|PE(O-18:1_22:6) | - | - | 1.598 | 1.82E-03 |
| 169 | PI(34:2)\|PI(16:0_18:2) | 1.683 | 1.97E-02 | 2.508 | 5.81E-06 |
| 170 | PI(36:2)\|PI(18:0_18:2) | - | - | 2.643 | 1.07E-05 |
| 171 | PI(36:3) | - | - | 1.783 | 1.01E-03 |
| 172 | PI(36:3)\|PI(18:1_18:2) | - | - | 2.957 | 3.17E-06 |
| 173 | PI(36:4)\|PI(16:0_20:4) | 1.463 | 1.53E-02 | 1.684 | 3.09E-03 |
| 174 | PI(37:4)\|PI(17:0_20:4) | - | - | 1.284 | 4.59E-02 |
| 175 | PI(38:5)\|PI(18:1_20:4) | 1.660 | 1.43E-02 | 2.605 | 1.93E-06 |

| | | | | | |
|---|---|---|---|---|---|
| 176 | PI(38:6)\|PI(16:0_22:6) | 2.383 | 2.99E-02 | 4.488 | 1.86E-05 |
| 177 | SM(32:1;2O) | 1.521 | 1.25E-02 | 2.324 | 1.42E-06 |
| 178 | SM(33:1;2O) | 1.816 | 5.03E-03 | 2.421 | 2.74E-06 |
| 179 | SM(34:1;2O) | 1.912 | 5.61E-03 | 3.225 | 7.48E-07 |
| 180 | SM(34:2;2O) | 2.941 | 3.12E-04 | 5.181 | 9.15E-09 |
| 181 | SM(35:1;2O) | 1.870 | 2.19E-02 | 2.839 | 2.27E-06 |
| 182 | SM(35:2;2O) | 1.725 | 4.62E-03 | 2.593 | 1.40E-06 |
| 183 | SM(36:1;2O) | 2.449 | 9.67E-03 | 6.149 | 1.09E-07 |
| 184 | SM(36:2;2O) | 1.821 | 1.09E-02 | 2.989 | 1.70E-06 |
| 185 | SM(36:3;2O) | - | - | 1.960 | 2.97E-04 |
| 186 | SM(37:1;2O) | - | - | 3.043 | 1.07E-05 |
| 187 | SM(38:1;2O) | - | - | 3.496 | 2.39E-06 |
| 188 | SM(38:2;2O) | - | - | 2.071 | 1.44E-03 |
| 189 | SM(39:1;2O) | - | - | 1.978 | 6.47E-04 |
| 190 | SM(40:1;2O) | - | - | 1.889 | 4.42E-04 |
| 191 | SM(40:2;2O) | 2.254 | 1.44E-02 | 4.148 | 6.84E-07 |
| 192 | SM(41:2;2O) | 1.544 | 2.18E-02 | 1.448 | 1.33E-02 |
| 193 | SM(42:1;2O) | - | - | 1.442 | 1.01E-02 |
| 194 | SM(42:2;2O) | 1.562 | 2.36E-02 | 2.396 | 3.80E-06 |
| 195 | SM(42:3;2O)_1 | - | - | 1.448 | 8.98E-03 |
| 196 | SM(42:3;2O)_2 | 1.574 | 4.12E-02 | 2.018 | 3.37E-04 |
| 197 | SM(42:5;2O) | 7.349 | 6.04E-04 | 18.331 | 8.16E-09 |
| 198 | SM(42:7;2O) | 2.800 | 9.67E-03 | 8.130 | 1.66E-06 |
| 199 | SM(43:1;2O) | - | - | 1.444 | 4.53E-02 |
| 200 | SM(43:2;2O) | 1.641 | 1.09E-02 | 1.947 | 4.17E-04 |
| 201 | SM(43:3;2O) | 1.623 | 7.71E-03 | 2.264 | 3.55E-06 |
| 202 | SM(44:1;2O) | 2.024 | 1.22E-02 | 2.454 | 6.02E-05 |
| 203 | SM(44:2;2O) | 2.018 | 1.22E-02 | 2.557 | 4.71E-05 |
| 204 | SM(44:3;2O) | - | - | 2.263 | 1.38E-04 |
| 205 | SM(44:5;2O) | 2.285 | 7.71E-03 | 3.465 | 2.39E-06 |
| 206 | SM(44:6;2O) | 2.263 | 2.02E-03 | 3.086 | 2.39E-06 |
| 207 | TG(47:0)\|TG(15:0_16:0_16:0) | 0.619 | 1.28E-02 | 0.424 | 1.77E-04 |
| 208 | TG(48:0)\|TG(16:0_16:0_16:0) | - | - | 0.615 | 1.37E-03 |
| 209 | TG(48:1)\|TG(16:0_16:0_16:1) | 0.623 | 1.59E-03 | 0.671 | 1.49E-02 |
| 210 | TG(48:3) | 0.360 | 1.19E-04 | 0.388 | 1.37E-05 |
| 211 | TG(49:0)\|TG(16:0_16:0_17:0) | 0.632 | 2.69E-02 | 0.333 | 2.24E-05 |

| | | | | | |
|---|---|---|---|---|---|
| 212 | TG(49:1)\|TG(15:0_16:0_18:1) | 0.676 | 3.70E-02 | 0.546 | 8.96E-04 |
| 213 | TG(50:0)\|TG(16:0_16:0_18:0) | - | - | 0.422 | 4.13E-04 |
| 214 | TG(50:3)\|TG(16:0_16:1_18:2) | 0.472 | 2.62E-03 | 0.484 | 1.36E-03 |
| 215 | TG(50:4)\|TG(16:0_16:0_18:4) | 0.575 | 1.05E-02 | 0.548 | 2.44E-03 |
| 216 | TG(50:4)\|TG(16:0_16:2_18:2) | 0.435 | 2.87E-03 | 0.308 | 2.98E-05 |
| 217 | TG(50:5)\|TG(14:0_18:2_18:3) | 0.254 | 6.83E-04 | 0.233 | 1.17E-04 |
| 218 | TG(51:0)\|TG(16:0_17:0_18:0) | - | - | 0.213 | 1.07E-05 |
| 219 | TG(51:1)\|TG(16:0_17:0_18:1) | - | - | 0.651 | 9.07E-03 |
| 220 | TG(51:3)\|TG(15:0_18:1_18:2) | 0.498 | 4.16E-03 | 0.522 | 4.23E-03 |
| 221 | TG(51:4)\|TG(15:0_18:2_18:2) | 0.547 | 1.64E-02 | 0.362 | 9.18E-05 |
| 222 | TG(52:2)\|TG(16:0_18:1_18:1) | - | - | 1.929 | 2.12E-04 |
| 223 | TG(52:4)\|TG(16:0_18:2_18:2) | - | - | 0.517 | 1.10E-02 |
| 224 | TG(52:5)\|TG(16:0_18:2_18:3) | 0.455 | 7.02E-03 | 0.453 | 5.47E-03 |
| 225 | TG(52:6)\|TG(14:0_18:2_20:4) | 0.314 | 5.19E-04 | 0.349 | 1.27E-04 |
| 226 | TG(52:6)\|TG(16:2_18:2_18:2) | 0.463 | 2.23E-02 | 0.477 | 2.17E-02 |
| 227 | TG(53:0)\|TG(16:0_18:0_19:0) | 0.557 | 2.22E-02 | 0.167 | 5.58E-06 |
| 228 | TG(53:1) | - | - | 0.325 | 1.60E-04 |
| 229 | TG(53:4)\|TG(16:0_17:0_20:4) | 0.597 | 1.88E-02 | 0.231 | 1.55E-06 |
| 230 | TG(53:4)\|TG(17:0_18:2_18:2) | 0.557 | 1.28E-02 | 0.509 | 3.09E-03 |
| 231 | TG(53:5)\|TG(17:1_18:2_18:2) | - | - | 0.431 | 4.96E-02 |
| 232 | TG(54:1)\|TG(16:0_18:0_20:1) | - | - | 0.360 | 2.26E-03 |
| 233 | TG(54:2)\|TG(18:0_18:1_18:1) | - | - | 1.736 | 1.06E-02 |
| 234 | TG(54:4)\|TG(16:0_18:0_20:4) | - | - | 0.313 | 4.67E-03 |
| 235 | TG(54:5)\|TG(18:1_18:2_18:2) | 0.628 | 3.25E-02 | - | - |
| 236 | TG(54:6)\|TG(16:0_18:2_20:4) | 0.520 | 2.89E-03 | 0.380 | 4.57E-04 |
| 237 | TG(54:6)\|TG(18:2_18:2_18:2) | 0.468 | 1.15E-02 | 0.374 | 8.96E-04 |
| 238 | TG(54:7)\|TG(16:0_18:2_20:5) | 0.202 | 5.83E-04 | 0.059 | 8.54E-08 |
| 239 | TG(54:7)\|TG(18:2_18:2_18:3)_1 | 0.209 | 6.83E-04 | 0.118 | 1.14E-07 |
| 240 | TG(54:7)\|TG(18:2_18:2_18:3)_2 | 0.262 | 1.70E-02 | 0.190 | 2.53E-03 |
| 241 | TG(55:1)\|TG(16:0_18:1_21:0) | - | - | 0.176 | 2.38E-04 |
| 242 | TG(55:2)\|TG(15:0_20:1_20:1) | - | - | 0.376 | 5.47E-04 |
| 243 | TG(55:3)\|TG(18:1_18:2_19:0) | - | - | 0.624 | 1.54E-02 |
| 244 | TG(55:4)\|TG(17:0_18:0_20:4) | - | - | 0.230 | 5.80E-07 |
| 245 | TG(56:1)\|TG(16:0_16:0_24:1) | - | - | 0.229 | 1.54E-04 |
| 246 | TG(56:2)\|TG(18:0_18:1_20:1) | - | - | 0.281 | 2.53E-03 |
| 247 | TG(56:4) | 0.690 | 2.87E-03 | - | - |

| | | | | | |
|---|---|---|---|---|---|
| 248 | TG(56:4)\|TG(18:0_18:0_20:4) | - | - | 0.299 | 1.23E-05 |
| 249 | TG(56:4)\|TG(18:1_18:1_20:2) | - | - | 0.590 | 3.09E-03 |
| 250 | TG(56:7)\|TG(16:0_18:1_22:6) | - | - | 0.510 | 6.32E-03 |
| 251 | TG(56:7)\|TG(18:1_18:2_20:4) | 0.596 | 2.11E-03 | 0.667 | 1.62E-02 |
| 252 | TG(56:8)\|TG(16:0_18:2_22:6) | 0.522 | 4.26E-03 | 0.193 | 3.18E-05 |
| 253 | TG(56:8)\|TG(18:2_18:2_20:4) | 0.501 | 7.84E-04 | 0.325 | 9.09E-05 |
| 254 | TG(56:9)\|TG(18:2_18:2_20:5) | 0.147 | 5.45E-05 | 0.133 | 3.38E-06 |
| 255 | TG(57:1)\|TG(16:0_18:1_23:0) | - | - | 0.197 | 5.26E-05 |
| 256 | TG(57:2)\|TG(16:0_18:2_23:0) | - | - | 0.280 | 3.30E-04 |
| 257 | TG(57:3)\|TG(18:1_19:1_20:1) | - | - | 0.347 | 8.01E-04 |
| 258 | TG(57:8)\|TG(17:0_20:4_20:4) | 0.439 | 6.83E-04 | 0.275 | 2.04E-06 |
| 259 | TG(58:1) | - | - | 0.230 | 2.53E-04 |
| 260 | TG(58:10)\|TG(16:0_20:4_22:6) | 0.567 | 4.35E-03 | 0.323 | 4.71E-05 |
| 261 | TG(58:10)\|TG(18:2_20:4_20:4) | 0.570 | 1.07E-03 | 0.516 | 1.12E-02 |
| 262 | TG(58:11)\|TG(18:2_20:4_20:5) | 0.117 | 2.96E-05 | 0.074 | 4.61E-07 |
| 263 | TG(58:12)\|TG(18:2_20:5_20:5) | 0.080 | 9.85E-05 | 0.031 | 9.15E-09 |
| 264 | TG(58:2)\|TG(16:0_18:1_24:1) | - | - | 0.287 | 2.07E-03 |
| 265 | TG(58:3)\|TG(16:0_18:2_24:1) | - | - | 0.315 | 5.10E-03 |
| 266 | TG(58:4)\|TG(18:2_20:1_20:1) | - | - | 0.297 | 7.56E-04 |
| 267 | TG(58:5)\|TG(18:0_20:1_20:4) | - | - | 0.376 | 4.90E-04 |
| 268 | TG(58:7)\|TG(18:0_18:1_22:6)_1 | 0.586 | 2.67E-02 | 0.471 | 1.77E-04 |
| 269 | TG(58:7)\|TG(18:0_18:1_22:6)_2 | 0.591 | 1.28E-02 | 0.624 | 1.02E-02 |
| 270 | TG(58:8)\|TG(18:1_18:1_22:6) | 0.545 | 1.77E-03 | 0.571 | 2.44E-03 |
| 271 | TG(58:9)\|TG(18:1_18:2_22:6) | 0.709 | 2.24E-02 | - | - |
| 272 | TG(59:1) | - | - | 0.162 | 6.58E-05 |
| 273 | TG(59:2)\|TG(18:1_18:1_23:0) | - | - | 0.189 | 1.95E-04 |
| 274 | TG(59:3)\|TG(18:2_20:1_21:0) | - | - | 0.245 | 4.50E-04 |
| 275 | TG(59:4)\|TG(18:2_18:2_23:0) | - | - | 0.512 | 1.01E-02 |
| 276 | TG(60:0)\|TG(18:0_20:0_22:0) | - | - | 0.168 | 5.27E-05 |
| 277 | TG(60:1)\|TG(18:0_20:1_22:0) | - | - | 0.174 | 9.50E-05 |
| 278 | TG(60:10)\|TG(18:1_20:3_22:6) | 0.700 | 3.68E-02 | - | - |
| 279 | TG(60:11)\|TG(18:2_20:4_22:5) | 0.398 | 9.78E-05 | 0.255 | 2.04E-06 |
| 280 | TG(60:12)\|TG(16:0_22:6_22:6) | - | - | 1.408 | 2.58E-02 |
| 281 | TG(60:12)\|TG(18:2_20:4_22:6) | 0.404 | 9.78E-05 | 0.287 | 2.97E-04 |
| 282 | TG(60:13)\|TG(18:2_20:5_22:6) | 0.115 | 2.96E-05 | 0.047 | 1.47E-07 |
| 283 | TG(60:2)\|TG(16:0_18:1_26:1) | - | - | 0.263 | 1.13E-03 |

| | | | | | |
|---|---|---|---|---|---|
| 284 | TG(60:3)\|TG(18:0_18:2_24:1) | - | - | 0.336 | 3.68E-03 |
| 285 | TG(60:4)\|TG(18:1_18:2_24:1) | - | - | 0.480 | 2.11E-02 |
| 286 | TG(60:5)\|TG(16:0_20:4_24:1) | - | - | 0.198 | 2.80E-04 |
| 287 | TG(60:8)\|TG(18:1_20:1_22:6) | 0.688 | 1.74E-02 | 0.493 | 1.07E-05 |
| 288 | TG(61:1)\|TG(18:0_20:1_23:0) | - | - | 0.144 | 5.01E-05 |
| 289 | TG(61:2)\|TG(18:1_19:0_24:1) | - | - | 0.157 | 8.96E-05 |
| 290 | TG(62:12)\|TG(18:2_22:5_22:5) | 0.753 | 2.84E-02 | 0.492 | 5.72E-03 |
| 291 | TG(62:13)\|TG(18:1_22:6_22:6) | - | - | 0.533 | 1.30E-02 |
| 292 | TG(62:13)\|TG(18:2_22:5_22:6) | 0.363 | 2.16E-04 | 0.161 | 1.37E-05 |
| 293 | TG(62:14) | 0.282 | 5.13E-04 | 0.105 | 1.17E-06 |
| 294 | TG(62:2)\|TG(18:1_20:0_24:1) | - | - | 0.185 | 4.56E-04 |
| 295 | TG(62:3)\|TG(18:0_18:2_26:1) | - | - | 0.253 | 1.40E-03 |
| 296 | TG(62:4)\|TG(18:1_18:2_26:1) | - | - | 0.339 | 3.00E-03 |
| 297 | TG(62:4)\|TG(18:2_18:2_26:0) | - | - | 0.128 | 3.72E-05 |
| 298 | TG(62:5)\|TG(16:0_20:4_26:1) | - | - | 0.184 | 1.82E-04 |
| 299 | TG(63:3)\|TG(18:2_20:1_25:0) | - | - | 0.182 | 1.28E-04 |
| 300 | TG(64:16)\|TG(20:4_22:6_22:6) | 0.384 | 2.78E-04 | 0.162 | 4.70E-06 |
| 301 | TG(64:2)\|TG(18:1_22:1_24:0) | - | - | 0.197 | 4.50E-04 |
| 302 | TG(64:3) | - | - | 0.208 | 6.15E-04 |
| 303 | TG(64:6)\|TG(18:2_20:4_26:0) | - | - | 0.197 | 1.60E-04 |
| 304 | TG(65:2)\|TG(21:0_22:1_22:1) | - | - | 0.111 | 3.18E-05 |
| 305 | TG(66:2)\|TG(18:0_18:1_30:1) | - | - | 0.213 | 1.60E-04 |
| 306 | TG(66:3)\|TG(16:0_18:2_32:1) | - | - | 0.280 | 1.33E-03 |
| 307 | TG(66:4)\|TG(18:2_24:1_24:1) | - | - | 0.250 | 7.44E-04 |
| 308 | TG(68:3)\|TG(18:1_18:1_32:1) | - | - | 0.309 | 2.55E-03 |
| 309 | TG(68:4)\|TG(18:2_18:2_32:0) | - | - | 0.320 | 2.16E-03 |
| 310 | TG(O-52:0)\|TG(O-20:0_16:0_16:0) | - | - | 0.551 | 2.65E-03 |
| 311 | TG(O-54:0)\|TG(O-22:0_16:0_16:0) | 0.591 | 1.10E-02 | 0.386 | 2.44E-04 |
| 312 | TG(O-54:1)\|TG(O-20:0_16:0_18:1) | - | - | 0.672 | 2.53E-02 |
| 313 | TG(O-56:1)\|TG(O-20:0_18:0_18:1) | - | - | 0.496 | 1.43E-03 |
| 314 | TG(O-56:3)\|TG(O-20:0_18:1_18:2) | - | - | 0.538 | 1.33E-02 |

Abbreviation: CAR, acylcarnitines; CE, cholesteryl esters; Cer, ceramides; DG, diacylglycerols; FA, fatty acids; LPC, lysophosphatidylcholines; LPE, lysophosphatidylethanolamines; LPE(O-), ether-linked lysophosphatidylethanolamines; PC, phosphatidylcholines; PC(O-), ether-linked phosphatidylcholines; PE, phosphatidylethanolamines; PE(O-), ether-linked phosphatidylethanolamines; PI, phosphatidylinositols;

SM, sphingomyelins; TG, triacylglycerols; TG(O-), ether-linked triacylglycerols; FC, fold change; FDR, false discovery rate.

**Supplementary Table S3. Differential lipids in liver of amiodarone-treated rats compared with the control group.**

| No | Differential lipids | 100 mg/kg vs. control | | 300 mg/kg vs. control | |
|---|---|---|---|---|---|
| | | FC | FDR | FC | FDR |
| 1 | BMP(36:3)\|BMP(18:1_18:2) | 6.198 | 5.87E-04 | 11.422 | 7.19E-10 |
| 2 | BMP(38:5)\|BMP(18:1_20:4) | 7.969 | 6.32E-04 | 17.806 | 1.61E-07 |
| 3 | BMP(38:6)\|BMP(18:2_20:4) | 6.852 | 6.29E-04 | 16.984 | 2.96E-11 |
| 4 | BMP(40:7)\|BMP(18:1_22:6) | 65.690 | 2.07E-03 | 240.630 | 4.75E-12 |
| 5 | BMP(40:8)\|BMP(18:2_22:6) | 8.003 | 4.41E-03 | 30.499 | 7.28E-09 |
| 6 | BMP(42:10)\|BMP(20:4_22:6) | 6.966 | 3.27E-03 | 24.221 | 5.12E-10 |
| 7 | BMP(42:11)\|BMP(20:5_22:6) | - | - | 4.010 | 1.43E-06 |
| 8 | BMP(44:11)\|BMP(22:5_22:6) | 13.662 | 2.98E-03 | 26.461 | 1.30E-07 |
| 9 | CAR(16:0) | 17.195 | 1.66E-02 | 40.357 | 2.58E-05 |
| 10 | CAR(18:0) | 3.712 | 4.20E-02 | 4.581 | 4.28E-04 |
| 11 | CAR(18:1) | 35.759 | 8.34E-03 | 100.300 | 5.55E-07 |
| 12 | CAR(20:0) | 4.364 | 6.81E-03 | 4.865 | 3.27E-05 |
| 13 | CAR(20:1) | 7.538 | 7.91E-03 | 10.836 | 6.60E-05 |
| 14 | CAR(20:4) | 8.391 | 4.78E-03 | 19.651 | 4.58E-06 |
| 15 | CAR(24:0) | 3.141 | 2.74E-03 | 4.040 | 1.43E-06 |
| 16 | CAR(24:1) | 3.486 | 2.07E-03 | 3.272 | 8.67E-05 |
| 17 | CAR(26:1) | 6.684 | 9.92E-04 | 8.071 | 1.17E-06 |
| 18 | CE(18:1) | - | - | 0.674 | 4.76E-02 |
| 19 | CE(18:2) | - | - | 0.578 | 4.29E-02 |
| 20 | CE(20:4) | 2.161 | 4.52E-03 | 2.429 | 6.01E-04 |
| 21 | CE(20:5) | - | - | 3.802 | 1.22E-04 |
| 22 | CE(22:6) | - | - | 1.988 | 2.42E-02 |
| 23 | Cer(33:1;2O)\|Cer(17:1;2O/16:0) | 1.580 | 4.85E-02 | - | - |
| 24 | Cer(34:1;2O)\|Cer(18:1;2O/16:0) | - | - | 1.861 | 1.30E-02 |
| 25 | Cer(34:2;2O) | 1.879 | 5.00E-02 | 2.327 | 1.34E-02 |
| 26 | Cer(35:1;2O)\|Cer(18:1;2O/17:0) | - | - | 1.431 | 4.67E-02 |
| 27 | Cer(36:1;2O)\|Cer(18:1;2O/18:0) | - | - | 1.849 | 1.85E-02 |
| 28 | Cer(38:1;2O)\|Cer(18:1;2O/20:0) | - | - | 1.590 | 4.28E-02 |
| 29 | Cer(40:1;2O)\|Cer(18:1;2O/22:0) | - | - | 1.775 | 1.72E-03 |
| 30 | Cer(40:2;2O)\|Cer(18:1;2O/22:1) | - | - | 1.828 | 8.55E-03 |
| 31 | Cer(42:1;2O)\|Cer(18:1;2O/24:0) | 1.393 | 1.17E-02 | 1.234 | 4.95E-02 |

| | | | | | |
|---|---|---|---|---|---|
| 32 | Cer(42:2;2O)\|Cer(18:1;2O/24:1) | 1.533 | 2.74E-03 | 1.457 | 1.02E-03 |
| 33 | Cer(44:1;2O)\|Cer(18:1;2O/26:0) | 2.097 | 7.43E-03 | 2.499 | 7.69E-03 |
| 34 | Cer(44:2;2O)\|Cer(18:1;2O/26:1) | 1.775 | 2.80E-03 | 1.639 | 1.42E-02 |
| 35 | CL(68:6)\|CL(34:3_34:3) | 0.573 | 2.77E-02 | 0.457 | 5.36E-03 |
| 36 | CL(70:4)\|CL(16:0_18:1_18:1_18:2) | 1.760 | 9.68E-03 | 1.622 | 3.44E-02 |
| 37 | CL(70:5)\|CL(16:0_18:2_18:1_18:2) | 1.593 | 7.91E-03 | 1.470 | 2.84E-02 |
| 38 | CL(70:6) | 0.699 | 7.55E-03 | 0.528 | 5.01E-04 |
| 39 | CL(70:7)\|CL(16:1_18:2_18:2_18:2) | 0.563 | 7.91E-03 | 0.424 | 8.07E-04 |
| 40 | CL(72:6)\|CL(18:1_18:2_18:1_18:2) | - | - | 0.715 | 3.16E-03 |
| 41 | CL(72:7)\|CL(36:3_36:4) | 2.202 | 7.91E-03 | 1.779 | 2.49E-02 |
| 42 | CL(72:8)\|CL(18:2_18:2_18:2_18:2) | 1.437 | 7.55E-03 | - | - |
| 43 | CL(72:8)\|CL(36:4_36:4) | 1.396 | 2.44E-02 | - | - |
| 44 | CL(74:10)\|CL(18:2_18:2_18:2_20:4) | 1.370 | 1.42E-02 | - | - |
| 45 | CL(74:7)\|CL(18:1_18:2_18:1_20:3) | - | - | 0.429 | 1.89E-04 |
| 46 | CL(74:8)\|CL(36:4_38:4) | - | - | 0.534 | 1.91E-03 |
| 47 | CL(74:9)\|CL(18:2_18:2_18:2_20:3) | - | - | 0.524 | 1.72E-03 |
| 48 | DG(32:0)\|DG(16:0_16:0) | - | - | 1.628 | 1.08E-02 |
| 49 | DG(34:0)\|DG(16:0_18:0) | - | - | 1.931 | 1.67E-02 |
| 50 | DG(34:1)\|DG(16:0_18:1) | - | - | 2.474 | 1.72E-03 |
| 51 | DG(34:2)\|DG(16:0_18:2) | 0.590 | 7.91E-03 | - | - |
| 52 | DG(35:2)\|DG(17:0_18:2) | - | - | 1.459 | 2.62E-02 |
| 53 | DG(36:1)\|DG(18:0_18:1) | - | - | 4.766 | 1.16E-03 |
| 54 | DG(36:2) | - | - | 4.529 | 1.42E-03 |
| 55 | DG(36:3)\|DG(18:1_18:2) | 0.654 | 3.19E-02 | 1.530 | 4.12E-02 |
| 56 | DG(36:4)\|DG(18:2_18:2) | 0.619 | 2.45E-02 | 0.599 | 1.79E-02 |
| 57 | DG(36:5)\|DG(18:2_18:3) | 0.388 | 9.92E-04 | 0.477 | 2.96E-04 |
| 58 | DG(38:2)\|DG(18:1_20:1) | - | - | 2.629 | 2.39E-03 |
| 59 | DG(38:2)\|DG(20:0_18:2) | - | - | 1.554 | 4.03E-02 |
| 60 | DG(38:4)\|DG(18:0_20:4) | - | - | 0.697 | 3.37E-02 |
| 61 | DG(40:4)\|DG(20:0_20:4) | - | - | 0.616 | 3.16E-03 |
| 62 | DG(40:7)\|DG(18:1_22:6) | - | - | 0.584 | 3.37E-02 |
| 63 | DG(40:8)\|DG(18:2_22:6) | 0.434 | 7.91E-03 | 0.246 | 1.34E-04 |
| 64 | DG(O-41:4)\|DG(O-21:0_20:4) | 1.542 | 5.81E-03 | 1.583 | 2.02E-02 |
| 65 | FA(14:0) | - | - | 0.606 | 7.83E-03 |
| 66 | FA(14:1) | - | - | 0.537 | 1.81E-03 |
| 67 | FA(16:1) | 0.522 | 3.32E-02 | - | - |

| | | | | | |
|---|---|---|---|---|---|
| 68 | FA(16:3) | 0.484 | 3.85E-03 | 0.665 | 4.46E-02 |
| 69 | FA(18:3) | 0.284 | 9.92E-04 | 0.503 | 1.85E-02 |
| 70 | FA(19:0) | - | - | 0.516 | 8.81E-03 |
| 71 | FA(20:2) | 0.503 | 4.46E-02 | - | - |
| 72 | FA(20:5) | 0.174 | 9.14E-04 | 0.093 | 7.55E-07 |
| 73 | FA(22:3) | 2.717 | 7.12E-03 | 4.764 | 2.47E-05 |
| 74 | FA(22:5) | 0.506 | 2.22E-02 | 0.408 | 1.82E-03 |
| 75 | FA(22:6) | 0.650 | 4.30E-02 | 0.520 | 9.03E-04 |
| 76 | FA(23:0) | - | - | 0.785 | 3.88E-02 |
| 77 | FA(24:6) | - | - | 2.491 | 5.16E-03 |
| 78 | LPC(16:1-SN2) | - | - | 0.235 | 2.36E-04 |
| 79 | LPC(17:0-SN1) | - | - | 0.728 | 3.49E-04 |
| 80 | LPC(18:1-SN1) | - | - | 1.710 | 3.15E-02 |
| 81 | LPC(18:1-SN2) | - | - | 2.954 | 3.35E-03 |
| 82 | LPC(19:0-SN1) | - | - | 0.607 | 6.23E-05 |
| 83 | LPC(20:0-SN1) | - | - | 0.708 | 7.72E-03 |
| 84 | LPC(20:1-SN1) | - | - | 0.572 | 1.36E-03 |
| 85 | LPC(20:2-SN2) | - | - | 0.407 | 2.37E-04 |
| 86 | LPC(20:4-SN1) | - | - | 0.374 | 3.43E-03 |
| 87 | LPC(22:6-SN2) | - | - | 0.311 | 1.30E-02 |
| 88 | LPC(24:0-SN1) | - | - | 4.282 | 1.91E-05 |
| 89 | LPE(16:0) | 1.329 | 4.09E-02 | - | - |
| 90 | LPE(18:1) | - | - | 1.549 | 7.73E-03 |
| 91 | NAE(19:0) | - | - | 0.086 | 3.27E-06 |
| 92 | NAE(19:1) | 0.528 | 4.59E-02 | 0.206 | 4.31E-04 |
| 93 | NAE(21:0) | - | - | 0.054 | 8.02E-06 |
| 94 | NAE(21:1) | - | - | 0.150 | 4.50E-06 |
| 95 | NAE(21:2) | 0.483 | 3.00E-02 | 0.069 | 1.14E-05 |
| 96 | PC(30:2) | - | - | 0.372 | 1.60E-02 |
| 97 | PC(32:2)\|PC(14:0_18:2) | - | - | 0.543 | 2.16E-02 |
| 98 | PC(32:3)_2 | - | - | 0.452 | 3.77E-02 |
| 99 | PC(33:1)\|PC(15:0_18:1) | 1.405 | 1.78E-02 | 1.502 | 4.31E-04 |
| 100 | PC(34:1) | 1.534 | 5.46E-03 | - | - |
| 101 | PC(34:1)\|PC(16:0_18:1) | 2.170 | 3.68E-03 | 2.212 | 2.44E-04 |
| 102 | PC(34:2)\|PC(16:0_18:2) | 1.476 | 8.80E-03 | - | - |
| 103 | PC(34:3)\|PC(16:0_18:3) | - | - | 5.029 | 9.25E-06 |

| | | | | | |
|---|---|---|---|---|---|
| 104 | PC(35:0) | - | - | 0.725 | 1.04E-02 |
| 105 | PC(35:1)\|PC(17:0_18:1) | 1.753 | 3.85E-03 | 1.974 | 4.77E-04 |
| 106 | PC(35:2)\|PC(17:0_18:2) | 1.429 | 2.13E-02 | - | - |
| 107 | PC(35:4) | - | - | 0.454 | 1.55E-03 |
| 108 | PC(36:0)\|PC(18:0_18:0) | - | - | 0.571 | 1.89E-04 |
| 109 | PC(36:1)\|PC(18:0_18:1) | 1.493 | 3.85E-03 | 1.730 | 2.14E-03 |
| 110 | PC(36:2)\|PC(18:0_18:2) | 1.443 | 2.19E-02 | - | - |
| 111 | PC(36:3)\|PC(16:0_20:3) | - | - | 1.953 | 6.82E-03 |
| 112 | PC(36:3)\|PC(18:1_18:2) | - | - | 2.289 | 2.01E-02 |
| 113 | PC(36:4)_1 | 1.399 | 1.86E-02 | 1.499 | 1.50E-02 |
| 114 | PC(36:4)_2 | - | - | 0.610 | 6.40E-04 |
| 115 | PC(36:4)\|PC(16:0_20:4) | 1.391 | 2.52E-02 | - | - |
| 116 | PC(36:5)\|PC(16:1_20:4) | - | - | 0.584 | 1.30E-02 |
| 117 | PC(36:6) | - | - | 1.843 | 1.06E-02 |
| 118 | PC(36:6)\|PC(14:0_22:6) | - | - | 0.548 | 1.41E-02 |
| 119 | PC(37:2)\|PC(19:0_18:2) | - | - | 0.690 | 8.76E-03 |
| 120 | PC(37:4)\|PC(17:0_20:4) | - | - | 0.759 | 6.16E-03 |
| 121 | PC(38:1)_1 | 1.459 | 4.78E-03 | 1.289 | 4.29E-02 |
| 122 | PC(38:2)\|PC(18:0_20:2) | - | - | 0.548 | 1.14E-04 |
| 123 | PC(38:5)_1 | 1.462 | 6.76E-03 | 1.263 | 2.80E-02 |
| 124 | PC(38:5)\|PC(18:1_20:4) | 1.462 | 7.04E-03 | - | - |
| 125 | PC(38:6)\|PC(16:0_22:6) | 1.481 | 5.63E-03 | - | - |
| 126 | PC(38:6)\|PC(18:2_20:4) | 1.648 | 2.07E-03 | 1.682 | 9.47E-04 |
| 127 | PC(39:4)\|PC(19:0_20:4) | - | - | 0.595 | 2.34E-03 |
| 128 | PC(39:5) | - | - | 0.519 | 3.48E-06 |
| 129 | PC(40:2) | - | - | 1.580 | 1.83E-02 |
| 130 | PC(40:4)\|PC(18:0_22:4) | 1.473 | 8.80E-03 | - | - |
| 131 | PC(40:5) | 1.322 | 7.42E-03 | - | - |
| 132 | PC(40:6) | - | - | 0.354 | 7.81E-05 |
| 133 | PC(40:6)\|PC(18:0_22:6) | 1.293 | 2.49E-02 | - | - |
| 134 | PC(40:7) | - | - | 2.965 | 7.66E-04 |
| 135 | PC(40:7)\|PC(18:1_22:6) | 1.557 | 4.44E-03 | 1.628 | 1.94E-03 |
| 136 | PC(40:8)\|PC(20:4_20:4) | 2.146 | 1.67E-03 | 2.320 | 3.99E-05 |
| 137 | PC(40:9)\|PC(20:4_20:5) | - | - | 2.947 | 1.42E-03 |
| 138 | PC(42:10)\|PC(20:4_22:6) | 4.473 | 2.80E-03 | 5.713 | 4.84E-05 |
| 139 | PC(42:3) | - | - | 1.642 | 1.45E-02 |

| | | | | | |
|---|---|---|---|---|---|
| 140 | PC(42:4) | 1.302 | 2.22E-02 | 1.411 | 1.07E-02 |
| 141 | PC(42:5) | 1.251 | 4.82E-02 | 1.381 | 4.20E-02 |
| 142 | PC(42:6) | - | - | 0.598 | 1.49E-03 |
| 143 | PC(42:8) | - | - | 0.670 | 2.01E-02 |
| 144 | PC(42:9) | 2.174 | 1.63E-03 | 2.330 | 1.03E-04 |
| 145 | PC(44:11) | 2.013 | 1.53E-02 | 1.609 | 4.67E-02 |
| 146 | PC(44:12) | 2.877 | 2.19E-02 | 3.161 | 1.06E-03 |
| 147 | PC(O-31:2) | 1.333 | 4.94E-02 | 1.803 | 5.86E-04 |
| 148 | PC(O-32:0)\|PC(O-16:0_16:0) | - | - | 5.050 | 6.16E-07 |
| 149 | PC(O-34:0)\|PC(O-16:0_18:0) | - | - | 4.014 | 2.33E-04 |
| 150 | PC(O-34:1)\|PC(O-18:1_16:0) | 4.904 | 1.94E-02 | 10.940 | 5.70E-08 |
| 151 | PC(O-34:2)\|PC(O-16:0_18:2) | 2.574 | 3.62E-02 | 4.784 | 3.83E-05 |
| 152 | PC(O-36:0) | - | - | 4.061 | 1.73E-07 |
| 153 | PC(O-36:1) | - | - | 4.834 | 1.30E-05 |
| 154 | PC(O-36:2)_1 | 2.515 | 1.84E-02 | 4.761 | 2.02E-06 |
| 155 | PC(O-36:2)_2 | 2.525 | 1.89E-03 | 2.715 | 2.91E-04 |
| 156 | PC(O-36:4)\|PC(O-16:0_20:4) | 3.097 | 3.37E-02 | 5.379 | 4.66E-07 |
| 157 | PC(O-36:5)\|PC(O-16:1_20:4) | 1.439 | 3.57E-02 | 1.464 | 4.57E-02 |
| 158 | PC(O-37:5) | - | - | 0.541 | 3.91E-02 |
| 159 | PC(O-37:7) | - | - | 0.642 | 9.03E-03 |
| 160 | PC(O-38:4) | 2.911 | 3.14E-02 | 3.811 | 3.48E-06 |
| 161 | PC(O-38:4)\|PC(O-18:0_20:4) | 3.183 | 1.18E-02 | 5.206 | 3.33E-07 |
| 162 | PC(O-38:5) | - | - | 1.645 | 2.66E-03 |
| 163 | PC(O-38:5)\|PC(O-18:1_20:4) | 3.733 | 8.80E-03 | 5.309 | 2.75E-07 |
| 164 | PC(O-38:6) | 1.672 | 6.54E-03 | 2.639 | 1.91E-03 |
| 165 | PC(O-39:1) | 1.389 | 6.01E-03 | 1.333 | 3.35E-03 |
| 166 | PC(O-39:2) | 0.292 | 1.64E-02 | 0.002 | 4.75E-12 |
| 167 | PC(O-40:4) | - | - | 6.371 | 5.25E-07 |
| 168 | PC(O-40:4)\|PC(O-20:0_20:4) | 4.412 | 1.48E-02 | 11.310 | 7.28E-09 |
| 169 | PC(O-40:5) | 2.557 | 1.31E-02 | 4.086 | 3.73E-07 |
| 170 | PC(O-40:6) | - | - | 2.899 | 2.59E-05 |
| 171 | PC(O-42:4) | 4.943 | 6.36E-03 | 11.761 | 5.77E-08 |
| 172 | PC(O-42:5) | 3.666 | 1.14E-02 | 8.204 | 1.25E-06 |
| 173 | PC(O-42:6) | 2.352 | 5.52E-03 | 4.178 | 4.39E-06 |
| 174 | PC(O-44:6) | 2.204 | 2.55E-02 | 4.406 | 3.27E-06 |
| 175 | PE(34:1)\|PE(16:0_18:1) | 2.116 | 2.24E-02 | 1.910 | 1.50E-03 |

| | | | | | |
|---|---|---|---|---|---|
| 176 | PE(34:2)\|PE(16:0_18:2) | 1.446 | 1.55E-02 | - | - |
| 177 | PE(35:1)\|PE(17:0_18:1) | 1.813 | 1.20E-02 | 1.905 | 5.37E-04 |
| 178 | PE(36:1)\|PE(18:0_18:1) | - | - | 2.876 | 2.16E-02 |
| 179 | PE(36:2)\|PE(18:0_18:2)_1 | 1.492 | 8.33E-03 | 1.302 | 1.17E-02 |
| 180 | PE(36:3)\|PE(18:1_18:2) | - | - | 2.070 | 9.97E-03 |
| 181 | PE(36:4)\|PE(16:0_20:4) | 1.610 | 1.72E-03 | - | - |
| 182 | PE(36:5) | 0.597 | 8.33E-03 | 0.303 | 1.29E-05 |
| 183 | PE(37:2)\|PE(19:0_18:2) | - | - | 0.592 | 1.14E-02 |
| 184 | PE(37:4)\|PE(17:0_20:4) | 1.429 | 2.07E-03 | - | - |
| 185 | PE(37:6) | - | - | 0.514 | 2.51E-02 |
| 186 | PE(38:1) | 1.345 | 1.25E-02 | - | - |
| 187 | PE(38:2)\|PE(18:0_20:2) | - | - | 0.703 | 2.07E-02 |
| 188 | PE(38:3) | 1.379 | 1.07E-02 | - | - |
| 189 | PE(38:3)\|PE(18:0_20:3) | - | - | 0.614 | 1.94E-02 |
| 190 | PE(38:4)\|PE(16:0_22:4) | 1.261 | 2.74E-02 | - | - |
| 191 | PE(38:4)\|PE(18:0_20:4) | 1.375 | 1.75E-02 | - | - |
| 192 | PE(38:5)\|PE(18:1_20:4) | 1.488 | 2.59E-03 | 1.470 | 1.02E-02 |
| 193 | PE(38:6) | 2.759 | 2.80E-03 | 3.270 | 4.58E-06 |
| 194 | PE(38:6)\|PE(16:0_22:6) | 1.652 | 1.04E-02 | - | - |
| 195 | PE(39:4)\|PE(19:0_20:4)_1 | - | - | 0.622 | 1.03E-02 |
| 196 | PE(39:4)\|PE(19:0_20:4)_2 | - | - | 0.566 | 4.51E-04 |
| 197 | PE(39:5)\|PE(17:0_22:5) | - | - | 0.620 | 2.25E-03 |
| 198 | PE(40:4)\|PE(18:0_22:4) | 1.323 | 1.15E-02 | - | - |
| 199 | PE(40:5)\|PE(18:0_22:5)_1 | - | - | 0.606 | 9.69E-03 |
| 200 | PE(40:6)\|PE(18:1_22:5) | - | - | 0.427 | 1.29E-05 |
| 201 | PE(40:7)\|PE(18:1_22:6) | 2.054 | 1.67E-03 | 2.019 | 1.41E-03 |
| 202 | PE(40:8)\|PE(18:2_22:6) | 1.940 | 3.61E-03 | 2.506 | 1.13E-04 |
| 203 | PE(O-36:2)\|PE(O-18:1_18:1) | - | - | 1.621 | 3.80E-02 |
| 204 | PE(O-36:5)\|PE(O-16:1_20:4) | 1.395 | 1.35E-02 | - | - |
| 205 | PE(O-38:5)\|PE(O-16:1_22:4) | 1.717 | 1.20E-02 | 1.784 | 2.37E-03 |
| 206 | PE(O-38:6)\|PE(O-18:2_20:4) | 1.330 | 3.84E-02 | 1.431 | 3.44E-02 |
| 207 | PE(O-38:7)\|PE(O-16:1_22:6) | - | - | 0.674 | 3.12E-02 |
| 208 | PE(O-40:5)\|PE(O-18:1_22:4) | - | - | 1.914 | 8.55E-03 |
| 209 | PE(O-40:5)\|PE(O-20:1_20:4) | 1.477 | 1.20E-02 | 2.039 | 5.37E-04 |
| 210 | PE(O-40:7)\|PE(O-18:1_22:6) | - | - | 0.642 | 2.01E-02 |
| 211 | PG(34:2)\|PG(16:0_18:2) | 3.447 | 5.52E-03 | 20.178 | 1.59E-05 |

| | | | | | |
|---|---|---|---|---|---|
| 212 | PG(34:2)\|PG(16:1_18:1) | 7.262 | 5.87E-04 | 15.534 | 6.66E-09 |
| 213 | PG(36:1)\|PG(18:0_18:1) | 1.928 | 4.78E-03 | 5.863 | 3.64E-03 |
| 214 | PG(36:2)\|PG(18:0_18:2) | 1.884 | 2.07E-03 | 11.809 | 5.85E-03 |
| 215 | PG(36:2)\|PG(18:1_18:1) | 6.368 | 7.45E-04 | 20.199 | 5.05E-07 |
| 216 | PG(36:3)\|PG(18:1_18:2)_1 | 3.841 | 6.32E-04 | 16.059 | 5.15E-06 |
| 217 | PG(36:3)\|PG(18:1_18:2)_2 | - | - | 19.087 | 1.17E-02 |
| 218 | PG(36:4)\|PG(16:0_20:4) | - | - | 5.130 | 8.77E-03 |
| 219 | PG(36:4)\|PG(18:2_18:2)_1 | 5.931 | 5.64E-04 | 12.075 | 3.60E-10 |
| 220 | PG(38:4)\|PG(18:0_20:4) | - | - | 7.898 | 3.61E-02 |
| 221 | PG(38:5)\|PG(18:1_20:4) | 5.456 | 1.27E-03 | 16.704 | 7.92E-09 |
| 222 | PG(38:7)\|PG(18:3_20:4) | 5.161 | 3.85E-03 | 15.419 | 3.69E-06 |
| 223 | PG(40:5)\|PG(18:1_22:4) | 5.259 | 5.87E-04 | 8.361 | 3.53E-07 |
| 224 | PG(40:7)\|PG(18:1_22:6) | 13.926 | 3.14E-03 | 54.481 | 3.13E-10 |
| 225 | PG(40:8)\|PG(18:2_22:6) | 10.514 | 4.41E-03 | 36.634 | 2.12E-10 |
| 226 | PG(40:9)\|PG(18:3_22:6) | 4.533 | 1.20E-02 | 16.705 | 2.13E-06 |
| 227 | PG(42:10)\|PG(20:4_22:6) | 8.370 | 4.88E-03 | 34.116 | 8.06E-10 |
| 228 | PG(42:11)\|PG(20:5_22:6) | - | - | 3.983 | 1.64E-06 |
| 229 | PG(44:11)\|PG(22:5_22:6) | 16.510 | 3.14E-03 | 36.769 | 1.05E-07 |
| 230 | PG(44:12)\|PG(22:6_22:6) | 4.863 | 1.82E-03 | 7.272 | 6.86E-07 |
| 231 | PI(34:1)\|PI(16:0_18:1) | 2.227 | 2.15E-02 | 2.704 | 1.64E-03 |
| 232 | PI(34:2)\|PI(16:0_18:2) | 2.694 | 5.28E-03 | 4.311 | 3.48E-05 |
| 233 | PI(35:2)\|PI(18:0_17:2) | 2.898 | 1.84E-02 | 3.582 | 1.77E-04 |
| 234 | PI(36:1)\|PI(18:0_18:1) | 2.674 | 3.27E-02 | 5.669 | 1.88E-04 |
| 235 | PI(36:2)\|PI(18:0_18:2) | 3.239 | 8.05E-03 | 5.501 | 1.63E-05 |
| 236 | PI(36:3)\|PI(18:1_18:2) | 2.558 | 9.47E-03 | 5.448 | 1.06E-04 |
| 237 | PI(36:4)\|PI(16:0_20:4) | 1.701 | 3.44E-02 | 2.609 | 2.64E-03 |
| 238 | PI(36:4)\|PI(18:2_18:2) | 2.857 | 8.34E-03 | 4.833 | 4.33E-04 |
| 239 | PI(38:4)\|PI(18:0_20:4) | 1.436 | 2.07E-03 | - | - |
| 240 | PI(38:5)\|PI(18:1_20:4) | 2.296 | 5.30E-03 | 4.396 | 8.95E-06 |
| 241 | PI(38:6)\|PI(16:0_22:6) | 3.168 | 1.82E-03 | 8.446 | 3.27E-06 |
| 242 | PI(40:5)\|PI(18:0_22:5) | - | - | 0.583 | 7.78E-03 |
| 243 | PS(36:4)\|PS(16:0_20:4) | 1.636 | 3.35E-03 | 2.118 | 3.25E-04 |
| 244 | PS(38:6)\|PS(16:0_22:6) | 2.347 | 2.59E-03 | 4.160 | 1.14E-05 |
| 245 | PS(40:6)\|PS(18:0_22:6) | 1.389 | 3.54E-02 | 1.436 | 1.33E-02 |
| 246 | SM(34:1;2O) | - | - | 1.608 | 1.06E-02 |
| 247 | SM(34:2;2O) | 1.598 | 2.01E-02 | 2.501 | 4.06E-04 |

| | | | | | |
|---|---|---|---|---|---|
| 248 | SM(36:1;2O) | 1.423 | 3.69E-02 | 1.691 | 2.54E-03 |
| 249 | SM(40:0;2O) | 0.785 | 4.16E-02 | - | - |
| 250 | SM(40:2;2O) | 1.629 | 3.60E-02 | 2.179 | 4.85E-04 |
| 251 | SM(41:1;2O) | 1.223 | 2.45E-02 | - | - |
| 252 | SM(41:8;2O) | - | - | 1.261 | 1.37E-02 |
| 253 | SM(41:9;2O) | - | - | 2.014 | 3.93E-03 |
| 254 | SM(42:1;2O)_2 | 1.253 | 1.15E-02 | - | - |
| 255 | SM(42:2;2O) | 1.439 | 2.20E-03 | 1.489 | 4.85E-04 |
| 256 | SM(42:3;2O) | 1.398 | 1.93E-02 | 1.746 | 6.45E-03 |
| 257 | SM(43:2;2O) | 1.353 | 3.44E-02 | - | - |
| 258 | SM(43:4;2O) | - | - | 0.675 | 4.31E-02 |
| 259 | SM(44:2;2O) | 1.944 | 7.45E-04 | 1.373 | 3.69E-02 |
| 260 | TG(48:0)\|TG(16:0_16:0_16:0) | - | - | 0.666 | 1.85E-02 |
| 261 | TG(48:1)\|TG(14:0_16:0_18:1) | 0.390 | 4.78E-03 | 0.515 | 2.23E-02 |
| 262 | TG(48:2)\|TG(14:0_16:0_18:2) | 0.268 | 1.82E-03 | 0.304 | 1.28E-03 |
| 263 | TG(48:3)\|TG(14:0_16:1_18:2) | 0.279 | 6.75E-03 | 0.214 | 4.85E-04 |
| 264 | TG(48:4)\|TG(12:0_18:2_18:2) | 0.246 | 2.07E-03 | 0.127 | 1.29E-05 |
| 265 | TG(49:1)\|TG(15:0_16:0_18:1) | 0.484 | 1.78E-02 | 0.586 | 4.28E-02 |
| 266 | TG(50:1)\|TG(16:0_16:0_18:1) | 0.521 | 1.25E-02 | - | - |
| 267 | TG(50:2)\|TG(16:0_16:0_18:2) | 0.444 | 7.06E-03 | 0.581 | 3.07E-02 |
| 268 | TG(50:3)\|TG(16:0_16:1_18:2) | 0.406 | 1.12E-02 | 0.422 | 8.32E-03 |
| 269 | TG(50:4)\|TG(14:0_18:2_18:2) | 0.339 | 6.75E-03 | 0.223 | 2.33E-04 |
| 270 | TG(51:1)\|TG(16:0_17:0_18:1) | 0.490 | 3.03E-02 | - | - |
| 271 | TG(51:2)\|TG(16:0_17:1_18:1) | 0.459 | 1.84E-02 | - | - |
| 272 | TG(51:3)\|TG(15:0_18:1_18:2) | 0.395 | 7.55E-03 | 0.311 | 1.46E-03 |
| 273 | TG(51:4)\|TG(15:0_18:2_18:2) | 0.339 | 6.81E-03 | 0.179 | 1.77E-04 |
| 274 | TG(52:1)\|TG(16:0_18:0_18:1) | 0.501 | 2.95E-02 | - | - |
| 275 | TG(52:3)\|TG(16:0_18:1_18:2) | 0.618 | 3.03E-02 | 0.662 | 3.89E-02 |
| 276 | TG(52:4)\|TG(16:0_18:2_18:2) | 0.491 | 1.34E-02 | 0.378 | 1.28E-03 |
| 277 | TG(52:4)\|TG(16:1_16:1_20:2) | 0.423 | 1.05E-02 | 0.243 | 3.67E-04 |
| 278 | TG(52:5)\|TG(16:0_18:2_18:3) | 0.352 | 8.34E-03 | 0.217 | 7.65E-04 |
| 279 | TG(52:6)\|TG(16:0_18:2_18:4) | 0.265 | 3.85E-03 | 0.118 | 9.53E-05 |
| 280 | TG(52:6)\|TG(16:2_18:2_18:2) | 0.219 | 3.85E-03 | 0.085 | 4.70E-05 |
| 281 | TG(52:7)\|TG(14:0_18:2_20:5) | 0.133 | 1.72E-03 | 0.042 | 2.31E-06 |
| 282 | TG(53:1)\|TG(17:0_18:0_18:1) | 0.525 | 1.85E-02 | 0.566 | 2.87E-02 |
| 283 | TG(53:2)\|TG(16:0_18:1_19:1) | 0.445 | 2.57E-02 | - | - |

| | | | | | |
|---|---|---|---|---|---|
| 284 | TG(53:3)\|TG(16:0_18:2_19:1) | 0.438 | 1.31E-02 | 0.417 | 1.30E-02 |
| 285 | TG(53:4)\|TG(17:0_18:2_18:2) | 0.393 | 8.76E-03 | 0.298 | 1.08E-03 |
| 286 | TG(53:5)\|TG(17:2_18:1_18:2) | 0.297 | 6.81E-03 | 0.189 | 2.75E-04 |
| 287 | TG(53:6)\|TG(15:0_18:2_20:4) | 0.369 | 8.53E-03 | 0.252 | 3.73E-04 |
| 288 | TG(54:2)\|TG(18:0_18:1_18:1) | 0.432 | 8.80E-03 | - | - |
| 289 | TG(54:3)\|TG(18:0_18:1_18:2) | 0.420 | 3.61E-03 | 0.675 | 2.93E-02 |
| 290 | TG(54:4)\|TG(18:1_18:1_18:2) | 0.369 | 3.85E-03 | 0.643 | 3.69E-02 |
| 291 | TG(54:5)\|TG(16:0_18:1_20:4) | 0.492 | 1.85E-02 | 0.495 | 8.77E-03 |
| 292 | TG(54:5)\|TG(18:1_18:2_18:2) | 0.320 | 4.18E-03 | 0.374 | 4.98E-03 |
| 293 | TG(54:6)\|TG(16:0_16:0_22:6) | 0.410 | 8.80E-03 | 0.331 | 1.24E-03 |
| 294 | TG(54:6)\|TG(16:0_18:2_20:4) | 0.425 | 1.14E-02 | 0.306 | 6.95E-04 |
| 295 | TG(54:6)\|TG(18:0_18:3_18:3) | 0.314 | 5.52E-03 | 0.158 | 1.38E-04 |
| 296 | TG(54:6)\|TG(18:2_18:2_18:2) | 0.313 | 8.80E-03 | 0.166 | 2.44E-04 |
| 297 | TG(54:7)\|TG(16:0_18:2_20:5) | 0.218 | 4.78E-03 | 0.060 | 5.18E-06 |
| 298 | TG(54:7)\|TG(18:2_18:2_18:3) | 0.187 | 4.25E-03 | 0.055 | 2.00E-05 |
| 299 | TG(54:8)\|TG(16:0_18:3_20:5) | 0.205 | 5.28E-03 | 0.075 | 1.44E-05 |
| 300 | TG(54:8)\|TG(16:1_18:2_20:5) | 0.112 | 1.87E-03 | 0.040 | 1.14E-05 |
| 301 | TG(55:2) | 0.573 | 2.74E-02 | 0.652 | 4.07E-02 |
| 302 | TG(55:3)\|TG(18:1_18:2_19:0) | 0.429 | 6.39E-03 | 0.638 | 3.46E-02 |
| 303 | TG(55:4)\|TG(18:2_18:2_19:0) | 0.390 | 7.34E-03 | 0.406 | 6.59E-03 |
| 304 | TG(55:5)\|TG(18:2_18:2_19:1) | 0.335 | 6.36E-03 | 0.276 | 1.01E-03 |
| 305 | TG(55:7)\|TG(17:1_18:2_20:4) | 0.282 | 1.20E-02 | 0.168 | 1.77E-04 |
| 306 | TG(55:8)\|TG(15:0_18:2_22:6) | 0.313 | 8.53E-03 | 0.141 | 5.21E-05 |
| 307 | TG(56:1)\|TG(18:0_18:1_20:0) | 0.500 | 2.15E-02 | 0.365 | 4.71E-03 |
| 308 | TG(56:3)\|TG(18:1_18:1_20:1) | 0.456 | 9.83E-03 | 0.659 | 3.37E-02 |
| 309 | TG(56:5) | 0.366 | 5.52E-03 | 0.848 | 2.43E-02 |
| 310 | TG(56:5)\|TG(16:0_18:1_22:4) | 0.504 | 3.66E-02 | 0.960 | 4.72E-02 |
| 311 | TG(56:6) | 0.414 | 1.42E-02 | 0.550 | 1.56E-02 |
| 312 | TG(56:7)\|TG(16:0_18:1_22:6) | 0.476 | 1.93E-02 | 0.336 | 1.08E-03 |
| 313 | TG(56:7)\|TG(16:0_18:2_22:5) | 0.326 | 7.04E-03 | 0.283 | 1.25E-03 |
| 314 | TG(56:8)\|TG(16:0_18:2_22:6) | 0.416 | 1.87E-02 | 0.198 | 2.06E-04 |
| 315 | TG(56:8)\|TG(18:2_18:2_20:4) | 0.277 | 6.81E-03 | 0.132 | 3.60E-05 |
| 316 | TG(58:1)\|TG(16:0_18:1_24:0) | - | - | 0.541 | 3.12E-02 |
| 317 | TG(58:10)\|TG(16:0_20:4_22:6) | 0.398 | 9.86E-03 | 0.247 | 2.38E-04 |
| 318 | TG(58:10)\|TG(18:2_18:2_22:6) | 0.229 | 3.68E-03 | 0.081 | 6.06E-06 |
| 319 | TG(58:12)\|TG(18:2_20:5_20:5) | 0.104 | 1.51E-03 | 0.104 | 1.42E-03 |

| | | | | | |
|---|---|---|---|---|---|
| 320 | TG(58:2)\|TG(22:0_18:1_18:1) | 0.542 | 2.13E-02 | 0.381 | 3.58E-03 |
| 321 | TG(58:3)\|TG(18:1_20:1_20:1) | 0.452 | 1.09E-02 | 0.357 | 2.24E-03 |
| 322 | TG(58:4)\|TG(18:1_20:1_20:2) | 0.586 | 3.18E-02 | 0.338 | 5.84E-04 |
| 323 | TG(58:6)\|TG(18:1_18:1_22:4) | 0.382 | 8.34E-03 | 0.449 | 5.16E-03 |
| 324 | TG(58:7)\|TG(18:0_18:1_22:6) | 0.351 | 3.14E-03 | 0.239 | 1.08E-04 |
| 325 | TG(58:7)\|TG(18:1_18:2_22:4) | 0.421 | 1.06E-02 | 0.481 | 5.60E-03 |
| 326 | TG(58:8)\|TG(18:1_18:1_22:6) | 0.256 | 2.36E-03 | 0.140 | 2.05E-05 |
| 327 | TG(58:9)\|TG(18:1_18:2_22:6) | 0.303 | 6.81E-03 | 0.169 | 1.38E-04 |
| 328 | TG(58:9)\|TG(18:2_18:2_22:5) | 0.212 | 5.38E-03 | 0.112 | 6.45E-05 |
| 329 | TG(60:10) | 0.430 | 1.35E-02 | 0.369 | 1.40E-03 |
| 330 | TG(60:11)\|TG(18:1_20:4_22:6) | 0.397 | 9.39E-03 | 0.220 | 1.67E-04 |
| 331 | TG(60:11)\|TG(18:2_20:4_22:5) | 0.264 | 7.25E-03 | 0.140 | 6.84E-05 |
| 332 | TG(60:12) | 0.177 | 6.39E-03 | 0.285 | 1.42E-02 |
| 333 | TG(60:12)\|TG(16:0_22:6_22:6) | 0.315 | 9.09E-03 | 0.091 | 2.99E-05 |
| 334 | TG(60:12)\|TG(18:2_20:4_22:6) | 0.210 | 2.74E-03 | 0.060 | 3.48E-06 |
| 335 | TG(60:2)\|TG(18:1_20:1_22:0) | - | - | 0.451 | 1.08E-02 |
| 336 | TG(60:3)\|TG(18:1_18:2_24:0) | 0.539 | 2.49E-02 | 0.369 | 1.80E-03 |
| 337 | TG(60:6)\|TG(18:1_20:1_22:4) | - | - | 0.547 | 2.42E-02 |
| 338 | TG(60:8)\|TG(18:0_20:4_22:4) | 0.448 | 1.32E-02 | 0.264 | 3.25E-04 |
| 339 | TG(62:11)\|TG(18:1_22:5_22:5) | - | - | 0.528 | 1.08E-02 |
| 340 | TG(62:12) | 0.504 | 3.00E-02 | 0.247 | 4.24E-04 |
| 341 | TG(62:12)\|TG(18:0_22:6_22:6) | 0.413 | 1.29E-02 | 0.139 | 3.48E-05 |
| 342 | TG(62:13) | 0.320 | 9.70E-03 | 0.081 | 4.06E-05 |
| 343 | TG(O-50:1)\|TG(O-18:1_16:0_16:0) | 0.435 | 1.48E-02 | 0.212 | 8.81E-05 |
| 344 | TG(O-52:1)\|TG(O-18:1_16:0_18:0) | 0.383 | 1.15E-02 | 0.184 | 1.26E-04 |
| 345 | TG(O-52:2)\|TG(O-16:0_18:1_18:1) | 0.431 | 8.53E-03 | 0.206 | 1.15E-05 |
| 346 | TG(O-54:1)\|TG(O-18:0_18:0_18:1) | - | - | 0.441 | 2.02E-02 |
| 347 | TG(O-54:2)\|TG(O-18:0_18:1_18:1) | 0.556 | 2.45E-02 | 0.389 | 2.13E-03 |

Abbreviations: BMP, bismonoacylglycerophosphates; CAR, acylcarnitines; CE, cholesteryl esters; Cer, ceramides; CL, cardiolipins; DG, diacylglycerols; DG(O-), ether-linked diacylglycerols; FA, fatty acids; LPC, lysophosphatidylcholines; LPE, lysophosphatidylethanolamines; NAE, N-acyl ethanolamines; PC, phosphatidylcholines; PC(O-), ether-linked phosphatidylcholines; PE, phosphatidylethanolamines; PE(O-), ether-linked phosphatidylethanolamines; PG, phosphatidylglycerols; PI, phosphatidylinositols; PS, phosphatidylserines; SM, sphingomyelins; TG, triacylglycerols; TG(O-), ether-linked triacylglycerols; FC, fold change; FDR, false discovery rate.
